# Viruses of Asgard archaea

**DOI:** 10.1101/2021.07.29.453957

**Authors:** Sofia Medvedeva, Jiarui Sun, Natalya Yutin, Eugene V. Koonin, Takuro Nunoura, Christian Rinke, Mart Krupovic

**Author notes:** Correspondence to Takuro Nunoura, Christian Rinke, Mart Krupovic. Evolutionary Biology of the Microbial Cell Unit, CNRS UMR2001, Department of Microbiology, Institut Pasteur, Paris, France.

## Abstract

Asgardarchaeota encode many eukaryotic signature proteins and are widely considered to represent the closest archaeal relatives of eukaryotes. Whether similarities between Asgard archaea and eukaryotes extend to their viromes remains unknown. Here we present 20 metagenome-assembled genomes of Asgardarchaeota from deep-sea sediments of the basin off the Shimokita Peninsula, Japan. By combining CRISPR spacer search of metagenomic sequences with phylogenomic analysis, we identify three family-level groups of viruses associated with Asgard archaea. The first group, Verdandiviruses, includes tailed viruses of the realm *Duplodnaviria*, the second one, Skuldviruses, consists of viruses with predicted icosahedral capsids that belong to the realm *Varidnaviria*, and the third group, Wyrdviruses, is related to spindle-shaped viruses previously identified in other archaea. More than 90% of the proteins encoded by these putative viruses of Asgard archaea show no sequence similarity to proteins encoded by other known viruses. Nevertheless, all three proposed families consist of viruses typical of prokaryotes, providing no indication of a specific evolutionary relationship between viruses infecting Asgard archaea and eukaryotes. Verdandiviruses and skuldviruses are likely to be lytic, whereas wyrdviruses, similar to all other known spindle-shaped viruses, probably establish chronic infection and are released without host cell lysis. All three groups of viruses were identified in sediment samples from distinct geographical locations and are expected to play important roles in controlling the Asgard archaea populations in deep-sea ecosystems.

## Text

The Asgard superphylum is an expansive group of metabolically versatile archaea that thrive primarily in anoxic sediments around the globe ^1–9^. Based on phylogenomic analyses, Asgard archaea have been originally classified into multiple phylum-level lineages, including Lokiarchaeota, Thorarchaeota, Odinarchaeota, Heimdallarchaeota, Helarchaeota, Sifarchaeota, Wukongarchaeota and several others, most of which were named after Norse gods ^1,5,7,9–12^. More recently, taxonomic rank normalization using relative evolutionary divergence has suggested that Asgard archaea represent a phylum, tentatively named Asgardarchaeota, including the classes Lokiarchaeia, Thorarchaeia, Odinarchaeia, Heimdallarchaeia, Sifarchaeia, Hermodarchaeia, Sifarchaeia, Baldrarchaeia, Wukongarchaeia and Jordarchaeia, with the other lineages classified as lower-rank taxa within the classes ^8,13^. The vast majority of these lineages have been discovered through metagenomics, whereas only one species has been isolated and successfully grown in the laboratory ^14^. Asgard archaea gained prominence due to their inferred key role in the origin of eukaryotes ^15^. Indeed, Heimdallarchaeia form a sister group to eukaryotes in most phylogenetic analyses ^1,5^, although alternative phylogenies have also been presented ^16,17^. Compared to other archaea, Asgard archaea encode a substantially expanded set of eukaryotic signature proteins (ESPs), including many proteins implicated in membrane trafficking, vesicle formation and transport, cytoskeleton formation, the ubiquitin network and other processes characteristic of eukaryotes ^1,5^. The ESPs are thought to have been inherited by the emerging eukaryotic cells from the last Asgard archaeal common ancestor. Should that be the case, a tantalizing question is whether eukaryotes also inherited viruses and other types of mobile genetic elements (MGE) from the Asgard archaea?

Viruses infecting archaea are notoriously diverse both in terms of their genome sequences and virion structures ^18–20^. Some archaeal viruses, in particular those with icosahedral virions, are evolutionarily related to bacterial and eukaryotic viruses, but the majority of archaeal virus groups are specific to archaea, with no identifiable relatives in the two other domains. Archaea-specific viruses often have odd-shaped virions, which resemble lemons, champagne bottles or droplets ^19^. Most archaeal viruses have been thus far isolated from hyperthermophilic or halophilic hosts, with only a handful of virus species described for methanogenic and ammonia-oxidizing archaea ^18^. No viruses infecting Asgard archaea have been isolated, primarily due to inherent difficulty to propagate Asgard hosts. Nevertheless, analysis of the CRISPR-Cas systems encoded by Asgard archaea revealed a remarkable diversity of defense systems in these organisms ^21^, implying a rich Asgard archaeal virome.

The functioning of the CRISPR-Cas system includes three major stages: (i) adaptation, where a Cas1-Cas2 complex, in some variants including additional Cas proteins, incorporates MGE-derived sequences (spacers) into a CRISPR array; (ii) processing, where the CRISPR array is transcribed and the transcript is processed into separate CRISPR (cr) RNAs containing the spacer sequence with 5′ and 3′ tags derived from the flanking repeats; (iii) interference, where the crRNA binds the CRISPR effector complex (also called interference module) which recognizes and degrades DNA and/or RNA of the cognate MGEs ^22,23^. CRISPR arrays are archives of past viral/MGE encounters, which can be harnessed to uncover the associations between viruses and their hosts. Indeed, matching the CRISPR spacers from a known organism to viruses, for which the identity of the host is not known, is widely used for host assignment for viruses discovered by metagenomics and arguably is the most straightforward and efficient approach to identify the hosts of such viruses ^24^.

Here we harness CRISPR spacer sequences from the sequenced Asgard archaea genomes to search for viruses infecting these organisms and describe three distinct family-level groups of putative Asgard viruses, all of which display typical features of viruses infecting bacteria or archaea.

## Results and Discussion

To search for viruses infecting Asgard archaea, we sequenced 12 metagenomes constructed from total environmental DNA directly extracted from the subseafloor sediments originating off the Shimokita Peninsula, Japan (site C9001, water depth 1,180◻m) ^25^ and representing different sediment depths ranging from 0.91 to 363.3 meters below the seafloor (mbsf). The sequence reads were assembled using MetaSpades ^26^ and MetaviralSPAdes ^27^ to obtain the metagenome-assembled genomes (MAGs) and genomes of viruses, respectively. Asgard archaeal MAGs were obtained from 7 of the 12 samples. A total of 20 Asgardarchaeota MAGs, including 12 Lokiarchaeia, 2 Thorarchaeia and 6 Heimdallarchaeia were analyzed in this study (Fig. 1), with an estimated completeness of 65.95 ± 17.93, estimated quality of 54.68 ± 15.78, and GC content of 32.78 ± 3.99 (Table S1).

**Figure 1.**
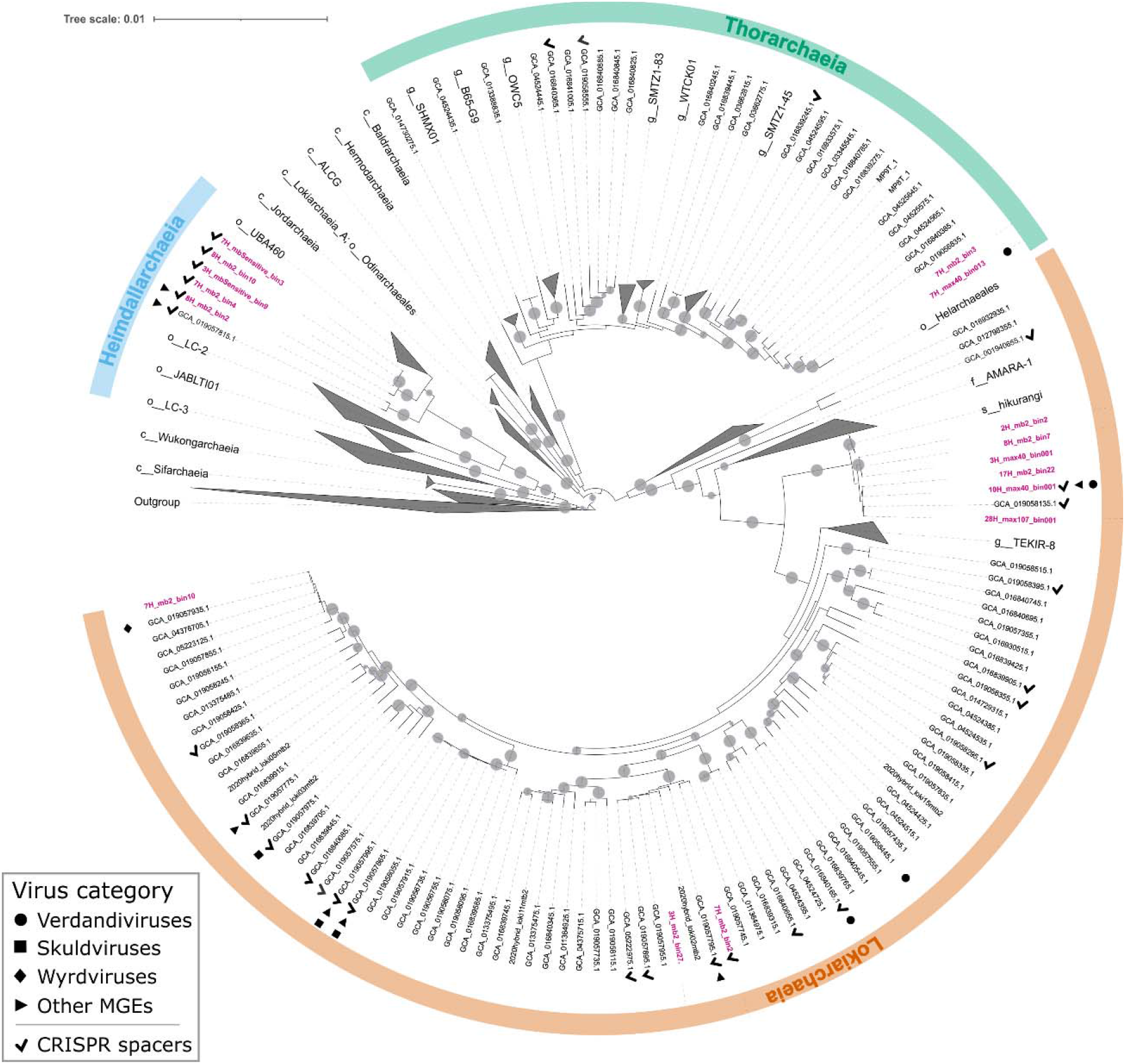
Phylogenomic tree of Asgardarchaeota. The alignment is based on a concatenated set of 53 protein markers (subsampled to 42 sites each, resulting in a total alignment length of 2058 sites) from 339 taxa. These taxa encompass 276 Asgardarchaeota MAGs, including 16 recovered in this study (highlighted in bold pink) (MAGs 3H_mb2_bin20, 4H_max40_bin02,1 4H_mb2_bin40, and 7H_mb2_bin7 were removed after the alignment trimming step due to low completeness), and 63 non-Asgardarchaeota species representatives from GTDB release RS95 as outgroup. Maximum-likelihood analysis was performed using IQ-TREE under the LG+C10+F+G+PMSF model. The tree is rooted on the non-Asgardarchaeota group. Nodes with bootstrap statistical support over 90% are indicated with grey dots. MAGs containing viral contigs and CRISPR arrays are indicated with different symbols and the key is provided in the bottom left corner.

To assign the putative viral genomes to Asgard archaeal hosts, we assembled a dataset of CRISPR spacers from Asgard archaeal MAGs obtained from the 20 Asgardarchaeota MAGs assembled in this study as well as from those reported previously (Fig. 2, Table S2). Analysis of the Asgard CRISPR arrays allowed us to define Asgard-specific CRISPR repeat sequences, which were used to identify additional CRISPR arrays from the contigs obtained from our sediment samples (Fig. 2A). In total, our CRISPR spacer dataset, originating from diverse geographical locations (Fig. 2B, Supplementary Data 1), included 2,545 spacers assigned to different Asgardarchaota lineages, with Lokiarchaeia contributing the highest number of spacers (Fig. 2C). All spacers were then used to search for protospacers in the 20 assembled MAGs and putative virus genomes from our dataset as well as contigs from GenBank and JGI sequence databases. In total, 14 contigs could be assigned to Asgard archaeal hosts based on CRISPR spacer matches (see Methods for details). Eight of the putative viral genomes originated from the metagenomes sequenced in this study (from depths ranging from 0.91 to 87.7 mbsf), whereas six additional genomes were recovered from datasets sequenced previously, including anoxic subseafloor sediment samples from two Pacific Ocean sites, the Hikurangi Subduction Margin (144.3 mbsf) ^8^ and Cascadia Margin (offshore of Oregon; (2 mbsf) ^28^, and one Indian Ocean site, Sumatra Forearc (1.6-6.07 mbsf; PRJNA367446). Each genome was targeted (≥90% identity between spacer and protospacer) by 1-6 CRISPR spacers assigned to putative Asgard classes Lokiarchaeia, Thorarchaeia, and Heimdalarchaeia (Fig. 1, Tables S3 and S4). Based on the conservation of viral hallmark proteins, including major capsid proteins (MCP), seven of the genomes were unequivocally identified as belonging to viruses from three unrelated groups, whereas the remaining seven could represent either novel viruses or other types of mobile genetic elements (MGEs; Fig. S1, see below). The viral dataset was further enriched by searching the JGI and GenBank sequence databases for related contigs. As a result, we collected 21 contig representing three groups of Asgard archaeal viruses originating from seven geographically remote locations (Fig. S2A). Network analysis of these viral genomes together with the bacterial and archaeal virus genomes available in RefSeq showed that the three Asgard virus groups are disconnected from each other as well as from other known viruses (Fig. S2B).

**Figure 2.**
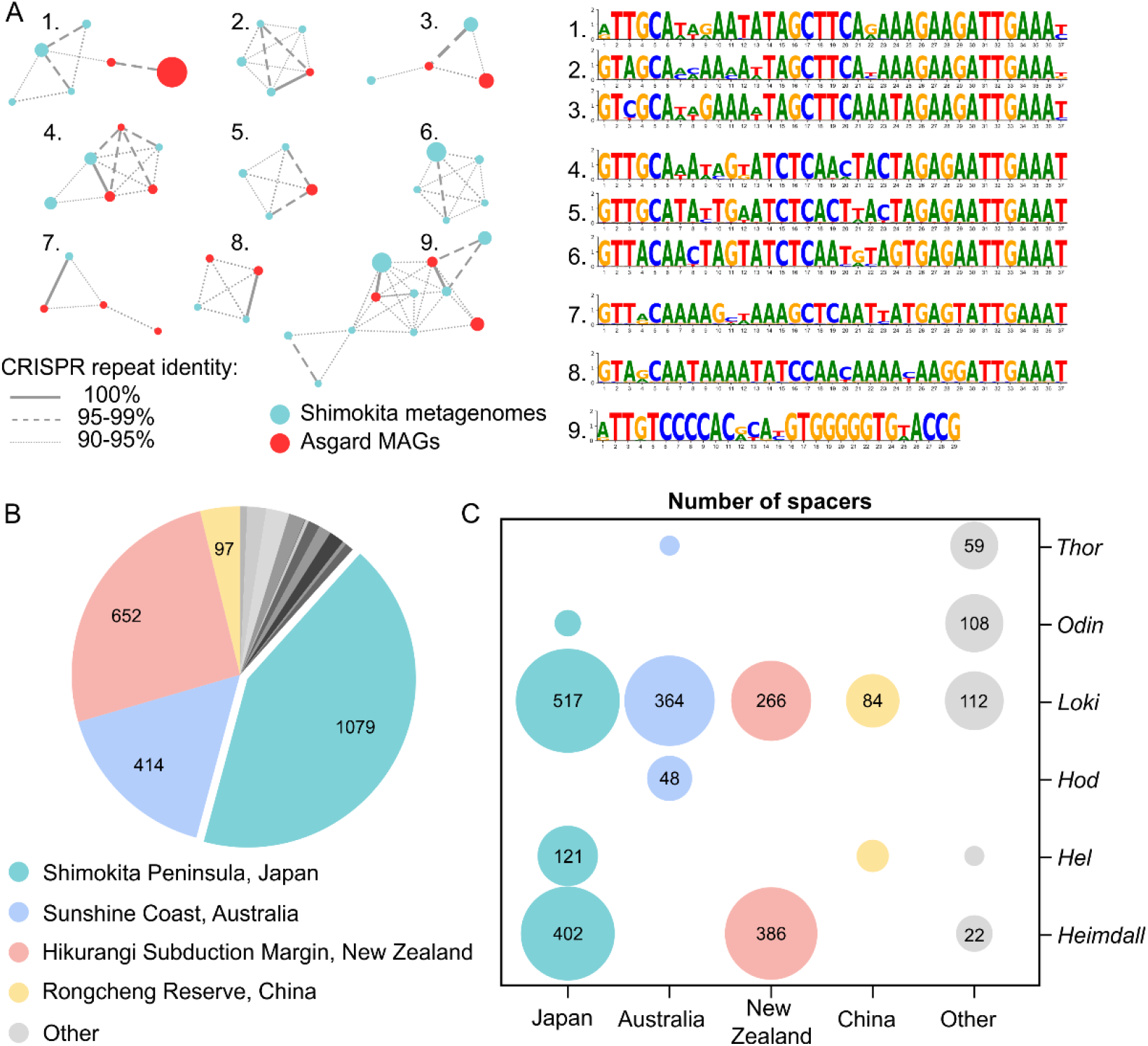
Description of the Asgardarchaeota CRISPR spacer dataset. A. Similarity of CRISPR repeats from metagenomic data (Bermuda) and CRISPR repeats from Asgardarchaeota MAGs (red). Unique CRISPR repeat sequences are shown as nodes in the network. The diameter of the node is proportional to the number of spacers associated with the repeat sequence in the dataset. Identical CRISPR repeats from metagenomic data and from Asgard MAGs are connected with solid lines. Dashed and dotted lines connect CRISPR repeat sequences with 95-99% and 90-95% identities, respectively. Consensus sequences for the nine major clusters are shown on the right. B. The source of the Asgardarchaeota CRISPR spacers. The number of spacers in the dataset originating from different geographical sites is indicated on the pie chart. C. Taxonomic distribution of Asgardarchaeota MAGs with CRISPR arrays for different isolation sites. The size of the circle is proportional to the number of spacers in the dataset.

### Verdandiviruses: Asgard archaeal tailed viruses of the class *Caudoviricetes*

Three of the genomes, VerdaV1, −2 and −3, assembled from the Shimokita dataset, encode the hallmark proteins specific to viruses of the class *Caudoviricetes*, which is currently the most widespread, environmentally abundant and genetically diverse group of viruses ^29,30^. Members of the *Caudoviricetes* infect bacteria and archaea, and have characteristic virions consisting of an icosahedral capsid and a helical tail attached to one of the capsid vertices. *Caudoviricetes* together with eukaryotic herpesviruses form the realm *Duplodnaviria* ^31^. Similar to previously characterized bacterial and archaeal viruses of the *Caudoviricetes*, each of the three virus genomes encodes the HK97-like major capsid protein (MCP), large subunit of the terminase (genome packaging ATPase-nuclease), the portal protein as well as several other structural proteins, including tail components (Fig. 3A). Structural modeling of the VerdaV1 MCP using RoseTTAFold, a recently introduced algorithm that outperforms all other currently publicly available structure prediction methods ^32^, yielded a model with the canonical HK97-like fold (Fig. 3B).

**Figure 3.**
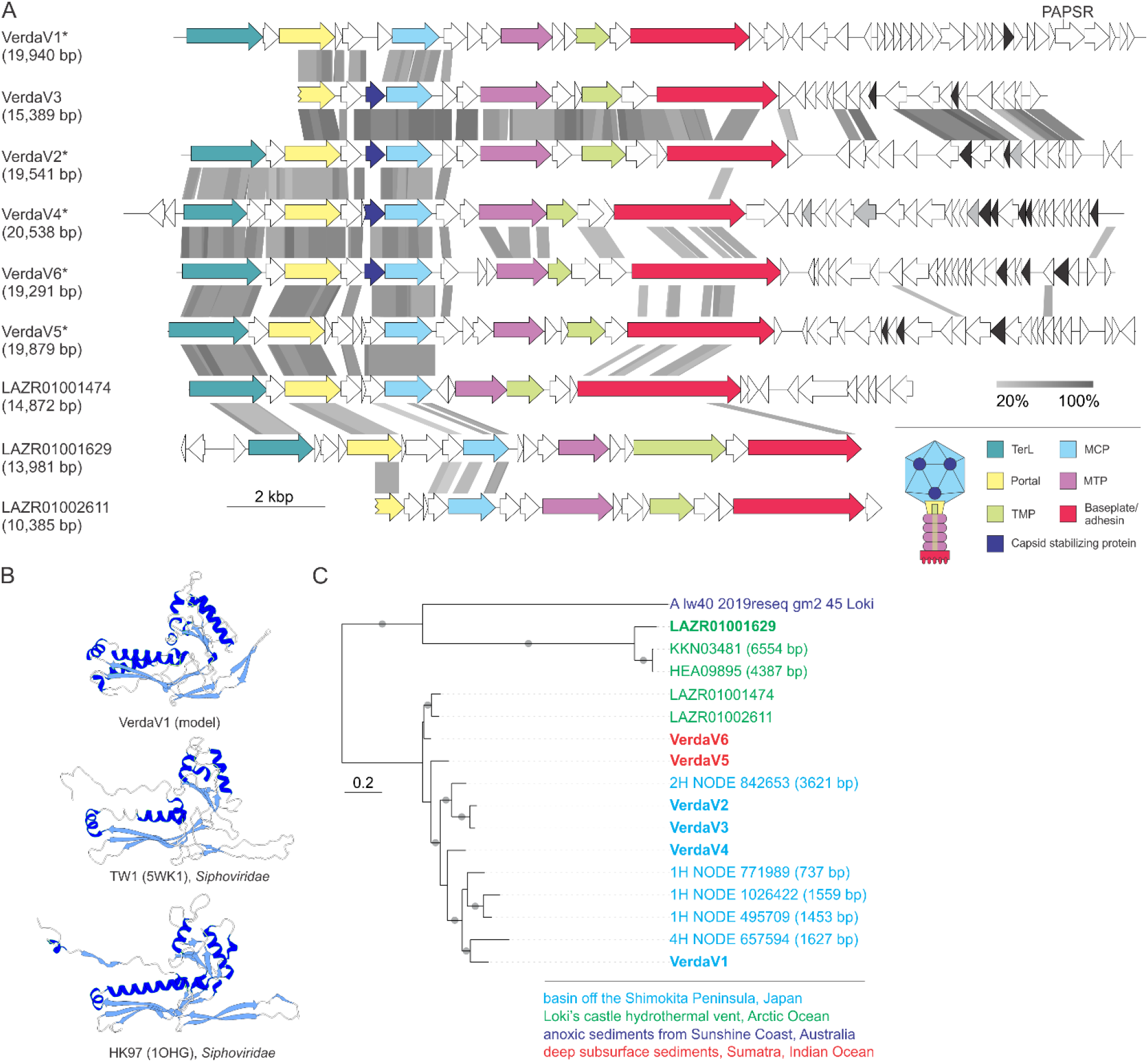
Diversity of verdandiviruses. A. Genome maps of verdnadiviruses. Homologous genes are shown using the same colors and the key is provided at the bottom of the panel. Also shown is the deduced schematic organization of the verdandivirus virion with colors marching those of the genes encoding the corresponding proteins. Genes encoding putative DNA-binding proteins with Zn-binding and helix-turn-helix domains are colored in black and grey, respectively. Grey shading connects genes displaying sequence similarity at the protein level, with the percent of sequence identity depicted with different shades of grey (see scale at the bottom). Asterisks denote complete genomes assembled as circular contigs. Abbreviations: PAPSR, phosphoadenosine phosphosulfate reductase; TMP, tail tape measure protein; MTP, major tail protein; MCP, major capsid protein; TerL, large subunit of the terminase. B. Comparison of the structural model of the major capsid protein of verdnadivirus AsTV-10H2 with the corresponding structures of siphoviruses TW1 and HK97. The models are colored according the secondary structure: α-helices, dark blue; β-strands, light blue. C. Maximul likelihood phylogeny of the major capsid proteins encoded by verdandiviruses. Taxa are colored based on the source of origin (key at the bottom of the figure), with those for which genomic contigs are shown in panel A shown in bold. The tree was constructed using the automatic optimal model selection (LG+G4) and is mid-point rooted. The scale bar represents the number of substitution per site. Circles at the nodes denote aLRT SH-like branch support values larger than 90%.

The VerdaV1 (19.9 kb) and VerdaV2 (19.5 kb) genomes were assembled as circular contigs (Table S3), suggesting that they correspond to complete, terminally redundant virus genomes, whereas VerdaV3 (15.4 kb) likely represents a partial genome (see below). Comparative genomics analysis showed that VerdaV1, −2 and −3 belong to the same virus group (Fig. 3A), which we refer to as ‘Verdandiviruses’ (for Verdandi, one of the three Norns, the most powerful beings in Norse mythology that govern the lives of gods and mortals). Given that the three genomes were identified in different sediment depths of offshore Shimokita Peninsula (at 18.7, 59.5 and 87.7 mbsf; Table S3), we addressed the possibility that related viruses could also be detected in samples from other depths. To this end, we performed BLASTP searches (E-value cutoff of 1e-05) queried with the corresponding MCP sequences against the assembled sequences from samples retrieved along the depth gradient. The analysis yielded six additional viral contigs and showed that related MCPs (and viruses) are also present in sediment samples from 0.9, 9.3 and 30.8 mbsf, indicating a broad distribution of verdandiviruses through the sediment column. One of the retrieved contigs (VerdaV4; 20.6 kb) was found to be circular and thus also corresponds to a complete virus genome. Furthermore, searches against the GenBank and JGI sequence databases yielded eight hits to virus-like contigs affiliated to Asgard archaea (Table S3). Five of the MCPs were encoded within large contigs (>10 kb) including two (Ga0114925_10000341 and Ga0114923_10001063 [VerdaV5 and VerdaV6, respectively]) circular ones. Notably, VerdaV5 was also targeted by a CRISPR spacer from a recently described Asgardarchaeota MAG ^1^, further supporting the host assignment for the Verdandivirus group, which is currently represented by five complete virus genomes (Fig. 3A, Table S3). Verdandivirus contigs were recovered from four geographically remote sampling sites (Fig. S2A) and maximum likelihood phylogenetic analysis of the verdandiviral MCPs revealed overall clustering according to the sampling site (Fig. 3C).

The vast majority of verdandiviral proteins (98%; 291 out of 298 proteins), including the MCP, display no recognizable similarity to proteins encoded by other known viruses (Table S5). Even the large subunit of the terminase (TerL), one of the most highly conserved proteins in *Caudoviricetes*, did not yield significant hits to the corresponding proteins of other bacterial or archaeal viruses, except in the case of LAZR01001629 (Table S5). Nevertheless, functional annotations were possible using sensitive profile-profile comparisons, in which HK97-like TerL and MCP were readily identified with highly significant scores (Fig. S3, Table S5). Verdandiviruses display considerable sequence conservation within the capsid formation and genome packaging modules (Fig. 3A). Notably, upstream of the MCP, all viruses carry a gene encoding a putative capsid stabilization protein distantly related to the corresponding protein of marine siphoviruses, e.g., TW1 ^33^, which might be important for maintaining capsid integrity under high hydrostatic pressures of the deep-sea ecosystems. Verdandiviruses encode tail proteins most closely similar to those of siphoviruses, including the major tail protein, tail tape measure protein and a baseplate hub protein with an adhesin domain, suggesting that verdandiviruses possess flexible, noncontractile tails. In most of these viruses, the tape measure protein is relatively short (median length 259 aa), suggesting that the tails themselves are not very long either. Assuming ~1.5 Å of tail length per amino acid residue of tape measure protein ^34,35^, the median tail length of verdandiviruses is predicted to be ~39 nm. The genes encoding tail proteins display low sequence conservation, whereas gene contents downstream of the tail modules are distinct in most of the identified viruses (Fig. 3A). Differences in the tail modules might correspond to different host ranges of the corresponding viruses. Indeed, whereas VerdaV1 and VerdaV5 were matched by CRISPR spacers from Lokiarchaeia, VerdaV2 and VerdaV3 are predicted to infect Thorarchaeia (Table S3). The distinct conservation patterns of the capsid and tail modules likely reflect modular evolution of these Asgard archaeal virus genomes, a common trait in bacterial and archaeal members of the *Caudoviricetes* ^36,37^.

Unlike many other members of *Caudoviricetes* with larger genomes, verdandiviruses do not encode any proteins implicated in virus genome replication or transcription, and thus, can be predicted to fully rely on the Asgard hosts for these processes. All complete verdandivirus genomes encompass arrays of short genes encoding predicted small, poorly conserved proteins, some of which are adjacent to genes encoding predicted helix-turn-helix transcriptional regulators. In many other bacterial and archaeal viruses, particularly among the *Caudoviricetes*, such arrays of small, fast evolving genes typically encode antidefense, and in particular, anti-CRISPR proteins (Acr) ^38,39^. Despite the lack of sequence similarity to known Acrs, the antidefense functions are likely to be represented in verdandiviruses as well.

### Skuldviruses: tailless icosahedral viruses of Asgard archaea

One of the contigs from Asgard archaeal MAGs assembled from anoxic sediments from the Hikurangi subduction margin ^8^, SkuldV1, targeted by two Asgard archaeal spacers (Table S3), encodes a double jelly-roll (DJR) MCP (Fig. 4A), a hallmark of viruses of the realm *Varidnaviria*, an expansive assemblage of viruses evolutionarily and structurally unrelated to duplodnaviruses ^31^. Similar to duplodnaviruses, varidnaviruses are environmentally abundant and infect hosts from all three domains of life ^31,40^. Sequence searches with the SkuldV1 MCP led to the identification of two additional contigs, one from our dataset (SkuldV2) and the other one from the GenBank database (JABUBK010000319, hereinafter referred to as SkuldV3). The latter contig was obtained from subseafloor sediments from the Cascadia Margin (Ocean Drilling Program, site 1244) in the Pacific Ocean ^28^ and is targeted by four spacers from our dataset. SkuldV1 and SkuldV3 were assembled as circular contigs and thus correspond to complete virus genomes, whereas SkuldV2 lacks the ~100 codons of the MCP gene (Fig. 4A). In addition to the DJR MCP, the vast majority of varidnaviruses encode genome packaging ATPases of the FtsK-HerA superfamily (unrelated to the terminase found in duplodnaviruses), which is responsible for pumping the viral DNA into icosahedral capsids through a conduit located at a unique vertex ^41^. A homolog of the genome packaging ATPase was identified in all three Asgard virus genomes, indicating that they are bona fide members of the realm *Varidnaviria*. We propose referring to this group of viruses as ‘Skuldviruses’ (for Skuld, another of the Norns).

**Figure 4.**
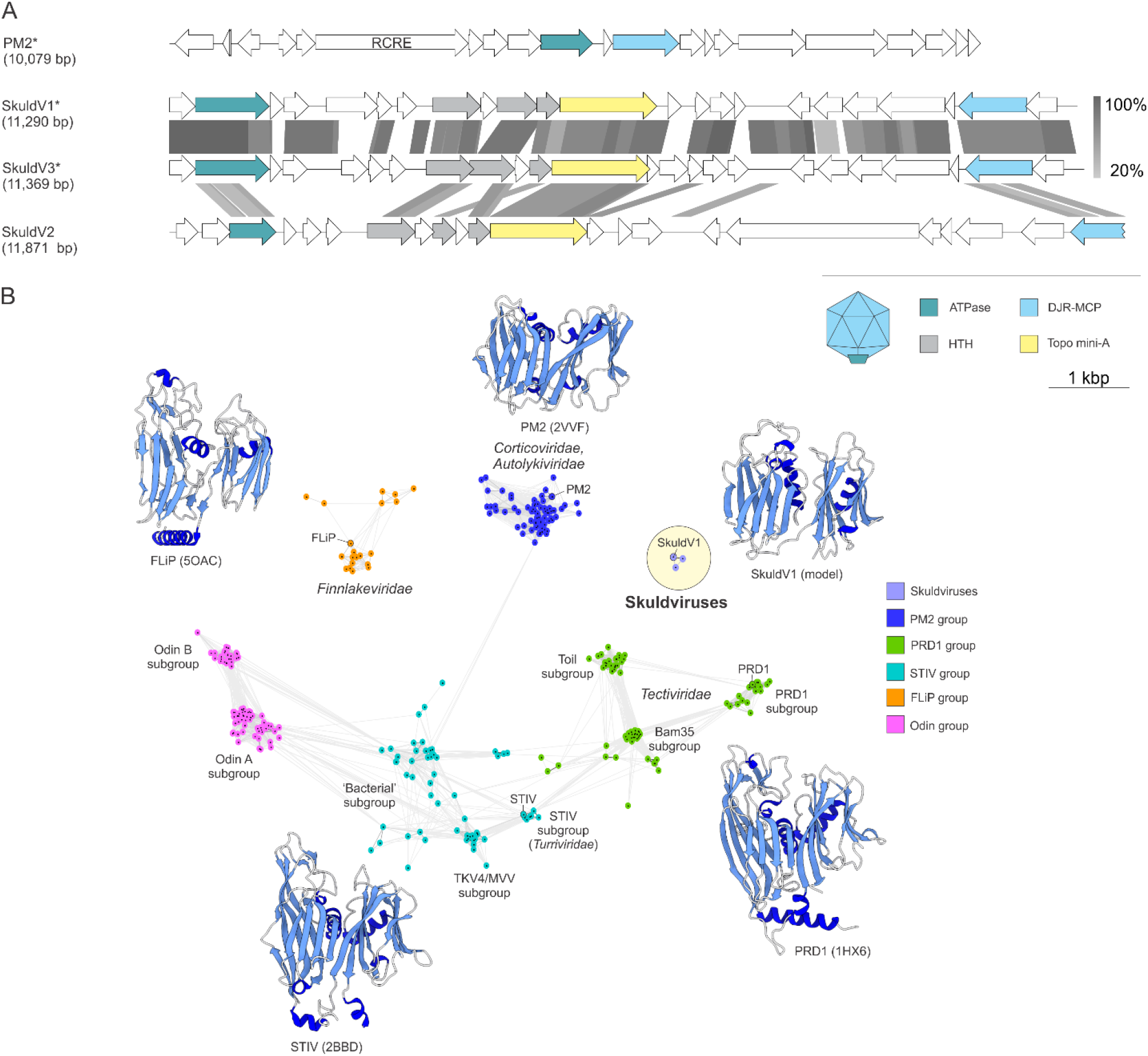
Diversity of skuldviruses. A. Genome maps of skuldviruses. Homologous genes are shown using the same colors and the key is provided at the bottom of the panel. Also shown is the deduced schematic organization of the skuldvirus virion with colors marching those of the genes encoding the corresponding proteins. Grey shading connects genes displaying sequence similarity at the protein level, with the percent of sequence identity depicted with different shades of grey (see scale on the right). Asterisks denote complete genomes assembled as circular contigs. Abbreviations: RCRE, rolling circle replication endonuclease; DJR-MCP, double jelly-roll major capsid protein; HTH, helix-turn-helix. B. Sequence similarity network of prokaryotic virus DJR MCPs. Protein sequences were clustered by the pairwise sequence similarity using CLANS. Lines connect sequences with CLANS P-value□≤□1e−04. Different previously defined groups ^43^ of DJR MCP are shown as clouds of differentially colored circles, with the key provided on the right. Note that the Odin group has been named as such previously because some of the MCPs in this cluster originated from the Odinarchaeia bin ^43^. PRD1, Toil, and Bam35 subgroups are named after the corresponding members of the family *Tectiviridae*. Skuldviruses are highlighted within a yellow circle. When available, MCP structures of viruses representing each group are shown next to the corresponding cluster (PDB accession numbers are given in the parenthesis). Skuldvirus cluster is represented by a structural model of the SkuldV1 MCP. The models are colored according the secondary structure: α-helices, dark blue; β-strands, light blue.

Similar to verdnadiviruses, the vast majority of skuldvirus proteins (97%; 66 out of 68 proteins) show no similarity to proteins encoded by other known viruses. Network analysis using CLANS ^42^ showed that the skuldvirus MCPs form a cluster separate from the previously characterized ^43^ groups of DJR MCPs of prokaryotic viruses (Fig. 4B). Consistently, BLASTP searches did not reveal homologs of skuldvirus MCPs in other known viruses. Nevertheless, profile-profile comparisons showed that it is most closely related to the corresponding proteins of prokaryotic viruses of the families *Corticoviridae* (bacteriophage PM2: HHsearch probability of 98.3), *Turriviridae* (archaeal virus STIV: HHsearch probability of 97.8) and *Tectiviridae* (bacteriophage PRD1: HHsearch probability of 96.2) (Fig. S4), whereas eukaryotic viruses with DJR MCPs were recovered with considerably lower scores (*Phycodnaviridae*, Pyramimonas orientalis virus: HHsearch probability of 83.4). Structural comparison of the SkuldV1 MCP model obtained using RoseTTAFold ^32^ (Fig. 4B) further showed that it is most similar to the MCP of Pseudoalteromonas phage PM2 (PDB id: 2VVF), a member of the family *Corticoviridae* ^44^, a group of viruses widespread in marine ecosystems ^45^. The MCP of corticoviruses is considered to possess a minimal DJR fold ^46^, lacking most of the structural embellishments present in the DJR MCPs of other viruses (Fig. 4B). Nevertheless, skuldviruses do not share genes with other known viruses other than those encoding the MCP and the genome packaging ATPase. Corticoviruses, turriviruses and tectiviruses, along with bacterial viruses of the family *Autolykiviridae* and eukaryotic *Adenoviridae* belong to the class *Tectiliviricetes* ^31^. We propose that skuldviruses represent a new virus family within a new order in the *Tectiliviricetes*.

Corticoviruses employ a rolling circle mechanism for genome replication and encode characteristic HUH superfamily endonucleases ^47^. No such genes or other putative replication genes were identified in skuldviruses. Instead, all three skuldviruses encode a protein related to the A subunit of type IIB topoisomerases, such as topoisomerase VI. The latter enzyme consists of two distinct subunits, A and B, with the catalytic tyrosine residue responsible for DNA nicking being located within the A subunit. Standalone A subunits, dubbed Topo mini-A, have been recently discovered in diverse bacterial and archaeal MGEs ^48^, but the functions of these proteins remain unknown. In the maximum likelihood phylogeny of Topo mini-A homologs, skuldvirus proteins form a clade with homologs from methanogenic Euryarchaeota (*Methanosarcina*) and ammonia-oxidizing Thaumarchaeota, which is nested between bacterial and archaeal homologs (Fig. S5). The skuldviral Topo mini-A might function as a replication protein, possibly initiating the rolling circle replication of the circular skuldvirus genomes, in a manner analogous to the HUH endonuclease. Obviously, this hypothesis awaits experimental validation.

Similar to verdandiviruses, scaldviruses encode two or three HTH proteins adjacent to small proteins without sequence similarity to any proteins in the current databases. These small proteins might function as Acrs.

### Wyrdviruses: Asgard viruses related to spindle-shaped archaeal viruses

One of the contigs from Asgard archaeal MAGs assembled from the Hikurangi Subduction Margin ^8^, WyrdV1 (15,570 bp), targeted by one CRISPR spacer from our collection, was found to encode two homologs of the MCPs specific to spindle-shaped archaeal viruses ^49^. In particular, the closest homolog was found in haloarchaeal virus His1 (family *Halspiviridae*) ^50^, whereas matches to the MCPs of evolutionarily related spindle-shaped viruses infecting hyperthermophilic Crenarchaeota (family *Fuselloviridae*) were recovered with lower scores (Fig. S6). His1-like MCPs are ~80 aa-long and contain two hydrophobic segments predicted to be membrane spanning domains ^49^. BLASTP searches using the WyrdV1 MCP as a query against the JGI and GenBank databases identified eight additional contigs longer than 10 kb (Fig. 5). All these contigs shared several genes encoding a morphogenetic module, including the MCP and receptor-binding adhesin, which in fuselloviruses and halspiviruses is located at one of the pointed ends of the virion ^51,52^. In addition, all these viruses encode a AAA+ ATPase, which in profile-profile comparisons showed the highest similarity to the morphogenesis (pI) proteins of bacterial filamentous phages (order *Tubulavirales*) ^53^. This ATPase is responsible for the extrusion of the viral genome through the cellular membrane during virion assembly ^54^. However, unlike pI-like proteins, but similar to the B251 ATPase of fuselloviruses, the ATPase of WyrdV1-like viruses lacks a transmembrane domain. A block of five genes, conserved in all WyrdV1-like viruses and located between the adhesin and the ATPase genes, is likely to be part of the morphogenetic module, although the encoded proteins are not detectably similar to proteins of any known viruses. We propose referring to this group of viruses as ‘Wyrdviruses’ (for Wyrd [Urðr], the third Norn). The homology between the tubulaviral and wyrdviral ATPases suggests that virion assembly of spindle-shaped Asgard archaeal viruses is mechanistically similar to the extrusion of filamentous bacteriophages. Indeed, spindle-shaped viruses infecting other archaea are released from the host without causing the cell lysis through a budding-like mechanism ^55,56^. Notably, some spindle-shaped viruses encode a single MCP (e.g., halspiviruses), whereas others encode two paralogous MCP (e.g., fuselloviruses). Similarly, wyrdviruses encode either one or two MCP paralogs (Fig. 5).

**Figure 5.**
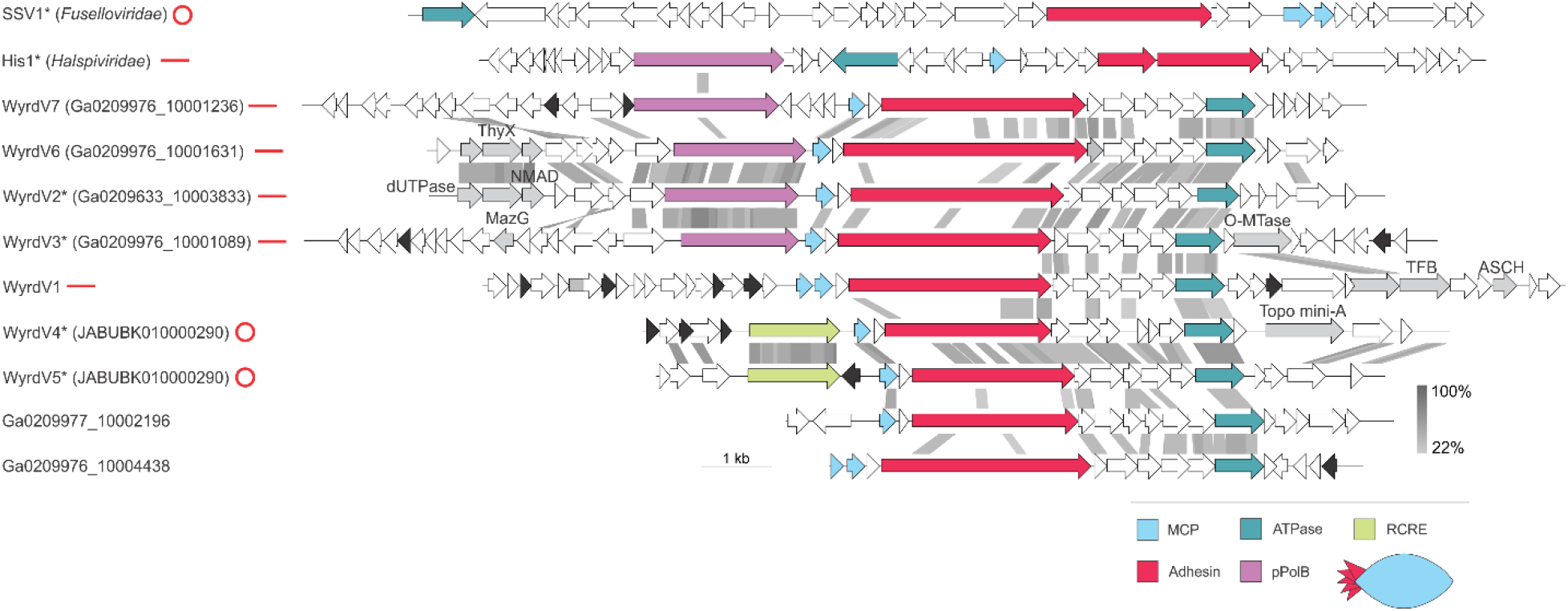
Diversity of wyrdviruses. Genome maps of wyrdviruses. Homologous genes are shown using the same colors and the key is provided at the bottom of the panel. Also shown is the deduced schematic organization of the skuldvirus virion with colors marching those of the genes encoding the corresponding proteins. Genes encoding putative DNA-binding proteins with Zn-binding and helix-turn-helix domains are colored in black and grey, respectively. Grey shading connects genes displaying sequence similarity at the protein level, with the percent of sequence identity depicted with different shades of grey (see scale on the right). Asterisks denote complete genomes assembled as circular contigs or linear contigs with terminal inverted repeats. The topology of each genome is shown next to the corresponding name: circle for circular genomes, lines for linear genomes. Abbreviations: ThyX, thymidylate synthase X; NMAD, nucleotide modification associated domain 1 protein; O-MTase, carbohydrate-specific methyltransferase; TFB, transcription initiation factor B; ASCH, ASC-1 homology domain; RCRE, rolling circle replication endonuclease; MCP, major capsid protein, pPolB, protein-primed family B DNA polymerase.

Similar to halspiviriruses, two of the contigs, Ga0209633_10003833 and Ga0209976_10001089, to which we refer to as WyrdV2 and WyrdV3, respectively, contained terminal inverted repeats (TIR), indicating that these are (nearly) complete genomes. By contrast, contigs JABUBK010000290 and JABUBK010000290, herein referred to as WyrdV4 and WyrdV5, respectively, contained direct terminal repeats, indicative of the completeness and circular structure of the genomes, resembling fuselloviruses. Consistent with the different genome structures, WyrdV2 and WyrdV3 (and halspiviruses) encode protein-primed family B DNA polymerases, whereas WyrdV4 and WyrdV5 encode rolling circle replication initiation endonucleases of the HUH superfamily (Fig. 5). Notably, WyrdV4, in addition, encodes Topo mini-A. The latter does not cluster with homologs from skuldviruses, but instead forms a clade with the Topo mini-A from the unassigned Asgard archaeal MGEs 10H_0 and 7H_42 (see below, Fig. S5). Contigs Ga0209976_10001631 (WyrdV6) and Ga0209976_10001236 (WyrdV7) also encode DNA polymerases and likely are linear, nearly complete viral genomes (Fig. 5). Remarkably, the DNA polymerase encoded by WyrdV7 is not orthologous to those of WyrdV2, WyrdV3 and WyrdV6 (Fig. 5). The latter group is most closely related (~30% identity) to the corresponding protein of an uncultured virus (MW522971) associated with Altiarchaeota ^57^, whereas the former protein is most similar to the DNA polymerase encoded by the spindle-shaped virus infecting marine thaumarchaea ^55^ (Table S3). Ga0209977_10002196 and Ga0209976_10004438 are partial genomes that lack the region encompassing the replication modules. Notably, WyrdV1 lacks either the polymerase or the rolling circle endonuclease gene and instead at the equivalent locus contains an array of short genes, some of which might encode replication initiators. Alternatively, however, these genes might function in antidefense or other aspects of virus-host interactions, whereas the replication machinery is fully provided by the host. Similar dramatic variation in gene content has been previously observed only in haloarchaeal viruses of the family *Pleolipoviridae*, where members of three genera encode non-homologous genome replication proteins (rolling circle endonucleases in alphapleolipoviruses and protein-primed family B DNA polymerases in gammapleolipoviruses) ^58^. Our present observations further illuminate the remarkable plasticity of the genome replication modules and genome structures in relatively closely related archaeal viruses, in general, and in wyrdviruses, in particular.

### Enigmatic MGEs of Asgard archaea

In addition to the three groups of viruses, we identified seven other contigs (7H_11, 8H_18, 8H_67, 10H_0, 7H_42, Ga0114923_10000127 and Ga0209976_10000148) targeted by Asgard archaeal CRISPR spacers (Table S3). Four of these contigs, 7H_11, 8H_18, 10H_0 and Ga0114923_10000127, were assembled as circular molecules and likely represent complete MGE genomes of 8,776 bp, 8,776 bp, 84,544 bp and 58,806 bp, respectively, whereas Ga0209976_10000148 (48,997 bp), 7H_42 (44,162 bp) and 8H_67 (13,282 bp) appear to be partial. The seven contigs do not encode identifiable homologs of major capsid proteins of known viruses and are likely to represent either novel viruses or non-viral MGEs, such as plasmids. Two circular contigs, 7H_11 and 8H_18, were nearly identical and targeted by identical spacers affiliated to Thorarchaeia (Tables S3 and S4), but were assembled from metagenomes originating from samples collected at different depths (59.5 and 68.8 mbsf, respectively). Notably, 7H_42 discovered in our samples from the offshore Shimokita Peninsula, Japan was found to be related to Ga0114923_10000127 and Ga0209976_10000148 originating from the sediment samples from the Sumatra, Indian Ocean (Fig. S1), attesting to the consistency of this emerging new group of Asgard MGEs.

The seven MGEs encode diverse proteins involved in DNA replication, repair and metabolism, which are common in MGE and viral genomes but, with the exception of 7H_42, Ga0114923_10000127 and Ga0209976_10000148 which form a group, display little overlap in gene content (see below). Nevertheless, 8H_18, 10H_0 and 7H_42 encode Topo mini-A homologs. Phylogenetic analysis of these proteins showed that, whereas 10H_0 and 7H_42 formed a clade with the Topo mini-A of wyrdvirus WyrdV4, 7H_42 branched among other archaeal sequences (Fig. S5), suggestive of active exchange of Topo mini-A genes among archaeal viruses and MGEs. The 10H_0 and 7H_42-like MGEs as well as verdandiviruses and wyrdviruses encode multiple non-orthologous Zn-finger proteins, which might be involved in transcription regulation or mediate protein-protein interactions. 10H_0 and 7H_42-like MGEs also share homologs of proliferating cellular nuclear antigen (PCNA) and transcription initiation factor B (TFB) (Fig. S1), both of which have been previously identified in archaeal viruses. For instance, PCNA is encoded by several tailed archaeal viruses infecting halophilic archaea ^59,60^ and spindle-shaped viruses of thaumarchaea ^55^, whereas TFB homologs are encoded by certain rod-shaped viruses infecting hyperthermophilic Crenarchaeota ^61^. 10H_0 also encodes Cdc6/Orc1-like origin recognition protein, nucleoside 2-deoxyribosyltransferase, DNA lyase, MazG-like nucleotide pyrophosphohydrolase and bifunctional (p)ppGpp synthase/hydrolase as well as several DNA methyltransferases and nucleases. By contrast, 7H_42-like MGEs encode a DNA primase-superfamily 3 helicase fusion protein that are commonly found in diverse MGEs including diverse varidnaviruses infecting eukaryotes, a Rad51-like recombinase, several nucleases and chromatin-associated proteins containing the HMG domain. The larger elements also encode auxiliary metabolic genes, including PAPS reductase, sulfatase, methylthiotransferase, and enzymes involved in carbohydrate metabolism, which could boost the metabolic activities of the respective hosts. For the smaller contigs, 8H_18 and 8H_67, the vast majority of genes were refractory to functional annotation even using the most sensitive available sequence similarity detection tools, such as HHpred ^62^.

### Auxiliary gene content of Asgard archaeal viruses

By dsDNA virus standards, genomes of verdandiviruses, skuldviruses and wyrdviruses are relatively small (≤20 kb). Thus, the corresponding gene contents are streamlined to include largely the core functions required for virion morphogenesis and genome replication. Nevertheless, some of these viruses encode auxiliary functions, including metabolic genes. In particular, verdandivirus VerdaV1 (and 10H_0 MGE) encode phosphoadenosine phosphosulfate (PAPS) reductase (also known as CysH), an enzyme reducing 3′– phosphoadenylylsulfate to phosphoadenosine-phosphate using thioredoxin as an electron donor. PAPS reductases have been previously identified in certain bacteriophages ^63–65^ and tailed haloarchaeal viruses ^59^, where they are thought to confer selective advantage to the host cells through facilitating sulfur metabolism and/or synthesis of sulfur-containing amino acids ^64^. PAPS reductase of VerdaV1 might perform a similar function.

Wyrdviruses WyrdV2 and WyrdV6 carry a block of three genes coding for dUTPase, thymidylate synthase X (ThyX) and an uncharacterized protein that is conserved in some phages and is annotated as nucleotide modification associated domain 1 protein (PF07659.13, DUF1599) (Fig. 5). This putative operon is likely to be involved in the biosynthesis of thymidylate from dUTP, to increase the pool of nucleotides available for the synthesis of viral DNA. WyrdV3 encodes a homolog of the nucleoside pyrophosphohydrolase MazG, which in bacteria prevents programmed cell death by degrading the central alarmone, ppGpp ^66^. MazG is highly conserved in tailed bacteriophages infecting cyanobacteria ^67^. Biochemical characterization of a cyanophage MazG has shown that, instead of degrading ppGpp, it preferentially hydrolyses dGTP and dCTP ^68^. Thus, MazG homolog in WyrdV3 might either function in disarming antiviral systems triggered by nucleotide-based alarmones, such as ppGpp, or in adjusting the intracellular nucleotide concentrations for optimal viral genome synthesis. Notably, MazG homologs are also encoded by Asgard archaeal MGEs 10H_0, 7H_42, Ga0114923_10000127 and Ga0209976_10000148 (Fig. S1).

None of the known archaeal viruses encodes its own RNA polymerase ^69^. Nevertheless, various transcription regulators with HTH, Zn-finger or ribbon-helix-helix domains are abundantly encoded in archaeal virus genomes. This is also the case with Asgard archaeal viruses described herein. Verdandiviruses and wyrdviruses encode multiple non-orthologous Zn-finger proteins, whereas skuldviruses encode several proteins with HTH domains (Fig. 4A). In addition, WyrdV1 (as well as 10H_0, 7H_42, Ga0114923_10000127 and Ga0209976_10000148) encodes a transcription initiation factor B (TFB), a homolog of eukaryotic TFIIB, which guides the initiation of RNA transcription ^70^. Among archaeal viruses, TFB homologs have been previously identified only in certain rod-shaped viruses infecting hyperthermophilic archaea ^61^. Thus, Asgard viruses appear to fully rely on the core transcription machinery of their hosts but encode various transcription factors that could be involved in the recruitment of this machinery for expression of viral genes as well as in the regulation of virus gene transcription. As mentioned above, some of the genes regulated by these transcription factors are likely to encode antidefense proteins.

WyrdV1 and WyrdV3 encode homologs of the carbohydrate-specific 3’-O-methyltransferase ^71^. In many archaeal viruses, the structural proteins are glycosylated by either the virus or host encoded glycosyltransferases, although the biological role of this post-translational modification remains unclear. The methyltransferase of WyrdV1 and WyrdV3 could participate in modification of the glycans attached to the virion proteins. Alternatively, the virus might alter the glycosylation makeup on the host cell surface as a strategy to avoid superinfection by other viruses.

### Concluding remarks: Evolutionary and ecological considerations

Here we describe three previously undetected, distinct groups of viruses associated with Asgard archaea of the lineages Lokiarchaeia and Thorarchaeia. Each group was identified in marine sediment samples from geographically remote sites (Fig. 1), suggesting a wide distribution of these viruses in Asgard archaea inhabited ecosystems. In addition, we recovered seven CRISPR-targeted MGEs associated with Lokiarchaeia, Thorarchaeia and Heimdallarchaeia that might represent distinct viruses with structural and morphogenetic proteins unrelated to those of any known viruses or, perhaps, more likely, plasmids. However, we envision that these seven MGEs will be characterized once cultures of the archaeal host lineages become available. Although verdandiviruses, skuldviruses and wyrdviruses, in all likelihood, do not comprise the complete Asgard virome, they provide important insights into the diversity and evolution of the Asgard viruses. All three virus groups are sufficiently distinct from previously characterized viruses to be considered as founding representatives of three new families. Wyrdviruses, inferred to form spindle-shaped virions, belong to one of the archaea-specific groups of viruses ^19^, thus far not observed in bacteria or eukaryotes. In archaea, spindle-shaped viruses are widely distributed and infect hyperthermophilic, halophilic and ammonia-oxidizing hosts from different phyla ^49,55^. Thus, wyrdviruses further expand the reach of spindle-shaped viruses to Asgard archaea, supporting the notion that this group of viruses was associated with the last archaeal common ancestor (LACA), and possibly, even the last universal cellular ancestor (LUCA) ^72^. By contrast, verdandiviruses and skuldviruses encode HK97-like and DJR MCPs and, at the highest taxonomic level, belong to the realms *Duplodnaviria* and *Varidnaviria*, respectively. Viruses of both realms have deep evolutionary origins and were proposed to have been present in both LUCA and LACA ^72^. The current work further supports this inference. Although members of the *Duplodnaviria* and *Varidnaviria* also infect eukaryotes, analyses of the verdandivirus and skuldvirus genome and protein sequences unequivocally show that they are more closely, even if distantly, related to their respective prokaryotic relatives. Thus, no putative direct ancestors of eukaryotic viruses were detected. Further exploration of the Asgard archaeal virome is needed to determine whether any of the virus groups associated with extant eukaryotes originate from Asgard viruses.

Due to inherent properties of the virion assembly and structure, all prokaryotic members of the realms *Duplodnaviria* and *Varidnaviria* are lytic viruses, which are released from the host cells by lysis (although some alternate lysis with lysogeny) ^73–75^. Hence, verdandiviruses and skuldviruses are also likely to kill their hosts at the end of the infection cycle, thereby promoting the turnover of Asgard archaea and nutrient cycling in deep-sea ecosystems. This possibility is consistent with previous results showing that viruses in deep-sea sediments lyse archaea faster compared to bacteria ^76^. By contrast, the mechanism of virion release employed by spindle-shaped viruses does not involve cell lysis, with virions continuously released from chronically infected cells ^55,56^. In the case of marine thaumarchaeal spindle-shaped virus NSV1, infection resulted in inhibition of the host growth and was accompanied by severe reduction in the rate of ammonia oxidation and nitrite reduction ^55^. Thus, infection dynamics and the impact of wyrdviruses is likely to be quite different compared to verdandiviruses and skuldviruses. Finally, some of the Asgard archaeal viruses carry auxiliary metabolic genes, such as the one encoding PAPS reductase, which might boost the metabolism of infected cells.

To conclude, the viral genomes described herein provide the first glimpse of the diversity and evolution of the Asgard archaeal virome and open the door for understanding virus-host interactions in the deep-sea ecosystems. Undoubtedly, many more Asgard archaeal virus groups remain to be discovered, which should clarify the contribution of these viruses to the evolution of the extant eukaryotic virome.

## Methods

### Site description and sampling

A total of 365 m of sediment cores were recovered from Hole C9001 C of Site C9001 (water depth of 1,180 m) located at the forearc basin off the Shimokita Peninsula, Japan (41°10.638N, 142°12.081E) by the drilling vessel Chikyu during JAMSTEC CK06-06 cruise in 2006. Coring procedure, subsampling for the molecular analyses, profiles of lithology, age model, porewater inorganic chemistry, organic chemistry and cell abundance, summary of the molecular microbiology in sediments at site C9001 were reported previously ^25,77^ (and references therein). Subsampled whole round cores (WRCs) for microbiology were stored at −80°C.

### Sample descriptions

A total of 12 sediment samples at depths of 0.9, 9.3, 18.5, 30.8, 48.3, 59.5, 68.8, 87.7, 116.4, 154.3, 254.7 and 363.3 mbsf were used in this study. These representatives were chosen based on porewater chemical profiles, lithostratigraphic properties, molecular biology data on the core sediments and F430 contents as follows. Note that 10 samples at depths of 0.9, 9.3, 18.5, 30.8, 48.3, 59.5, 116.4, 154.3, 254.3 and 363.3 mbsf were previously analyzed by SSU rRNA gene tag sequencing with a 454 FLX Titanium sequencer, and details have been reported in ref ^25^.

At 0.9 mbsf, the uppermost section of the core column was chosen. At 9.3 mbsf, the region just beneath the SMTZ was chosen. At 18.5 mbsf, highest relative abundance of Lokiarchaeia was found in the previous SSU rRNA gene tag sequencing. At 48.3 mbsf, relatively higher abundance of SAGMEG in the archaeal community was observed. At 59.5 mbsf predominance of Woesearchaeota was observed. For the sediment, at depth of 68.8 mbsf, the sample harbored anomalously high F430 concentrations, and the sample from a depth of 87.7 mbsf was used as a reference. Sediment at 116.4 mbsf consisted of ash/pumice, whereas other samples used in this study were pelagic clay sediments, and predominance of Bathyarchaeota was observed. Samples from 154.3, 254.3 and 363.3 mbsf were selected in 100 m depth intervals from the bottom of this drilling hole ^25,77^.

### DNA extraction, shotgun metagenome sequencing and SSU rRNA gene tag sequencing

DNA extraction and shotgun metagenome library construction were described in ref ^78^. Briefly, total environmental DNA was extracted from approximately 5 g of sediment subsampled from inner part of WRC using DNeasy PowerMax Soil Kit (Qiagen) with a minor modification ^79^. Then, further purification was performed, NucleoSpin gDNA Clean-up (MACHERY-NAGEL, Germany). The purified DNA was used for Illumina sequencing library construction using a KAPA Hyper Prep Kit (for Illumina) (KAPA Biosystems, Wilmington, MA, USA). Sequencing was performed on an Illumina HiSeq 2500 platform (San Diego, CA, USA) and 250-bp paired-end reads were generated.

### Assembly, binning, and Asgardarchaeota phylogeny

Contigs from each sample were assembled using MetaSpades v0.7.12-r1039 ^26^ and binned with UniteM (https://github.com/dparks1134/unitem). Metagenome-assembled genome (MAG) sets were generated after a dereplication of bins using DAS_Tool v1.1.2 ^80^ and quality check using CheckM ^81^. MAGs with an estimated quality (completeness - 4*contamination) of less than 30% were excluded in the downstream analysis. Taxonomic assignments of all MAGs were performed using the ‘classify’ function in GTDBtk ^82^.

The phylogenetic analysis included 20 Asgardarchaeota MAGs, together with 255 published Asgardarchaeota and 64 non-Asgardarchaeota archaeal genomes. A marker set consisting of 53 ribosomal proteins of the 239 genomes were identified and independently aligned using the ‘identify’ and ‘align’ functions in GTDBtk ^82^. The individual multiple sequence alignments were then concatenated and trimmed in GTDBtk using ‘trim_msa’ (--min_perc_aa 0.4). Maximum likelihood phylogenies of Asgardarchaeota MAGs were initially estimated by FastTree v2.1.11 ^83^ with default settings, and subsequently inferred using IQ-Tree v1.6.984 ^84^ under the LG+C10+F+G+PMSF model with 100 bootstraps. The final consensus tree was visualized and beautified in iTOL ^85^.

### Assembly and analysis of viral contigs

Potential viral and MGE contigs were assembled from metagenomic reads by MetaViralSPAdes pipeline with default parameters ^27^. Direct and inverted terminal repeats were identified by BLASTN ^86^. Open reading frames were identified with Prokka ^87^ and annotated using HHsearch ^88^ against Pfam, PDB, SCOPe, CDD and viral protein sequence databases. Sequences of major capsid proteins were used for BLASTP searches (E-value cutoff of 1e-05, 70% query coverage) against GenBank and JGI databases. Genomes of assembled viral genomes were compared and visualized using EasyFig ^89^ with the tBLASTx option. Network analysis of viral genomes was performed using vConTACT v2.0.9.19 with default parameters against Viral RefSeq v201 reference database ^90^. The virus network was visualized with Cytoscape ^91^.

### Collection of the CRISPR spacer dataset

CRISPR arrays were detected in metagenomic contigs and 255 published Asgardarchaeota MAGs using minced v0.4.2 ^92^. CRISPR arrays from metagenomic contigs were assigned to “Asgard” and “non-Asgard” groups based on CRISPR repeat similarity to previously characterized Asgard CRISPR repeats ^21^. CRISPR repeat sequences from metagenomic contigs and from Asgard MAGs were then clustered together using all-against-all BLASTN search (E-value cutoff of 1e-05, 90% identity, word size 7) and the result of clustering was visualized in Cytoscape ^91^. Consensus sequences for the 9 major clusters of CRISPR repeats were constructed using python package Logomaker ^93^. Spacer sequences from Asgard CRISPR arrays were matched to metagenomic contigs and published Asgard MAGs (BLASTN, E-value 1e-05, 90% identity, word size 7), and viral genomes available at IGV database (default parameters) at https://img.jgi.doe.gov/cgi-bin/m/main.cgi?section=WorkspaceBlast&page=viralform.

### Structural modeling and network analysis of the major capsid proteins

Structural modeling of the representative verdnadivirus and skuldvirus major capsid proteins was performed with RoseTTAFold ^32^. The MCPs of skuldviruses were compared to the DJR MCPs of other known prokaryotic viruses using CLANS ^42^, an implementation of the Fruchterman-Reingold force-directed layout algorithm, which treats protein sequences as point masses in a virtual multidimensional space, in which they attract or repel each other based on the strength of their pairwise similarities (CLANS p-values). The reference dataset of DJR MCP was obtained from Ref ^43^. Sequences were clustered using CLANS with BLASTP option (E-value of 1e-04) ^42^.

### Phylogenetic analysis of viral proteins

Sequences were aligned using MAFFT in ‘Auto’ mode ^94^. For phylogenetic analysis, uninformative positions we removed using TrimAl with gap threshold of 0.2 for the verdandiviral MCPs and gappyout option for Topo mini-A ^95^. The final alignments contained 323 and 277 positions, respectively. The maximum likelihood trees was inferred using IQ-TREE v2 ^84^. The best-fitting substitution models were selected by IQ-TREE and were LG+G4 and LG+R5 for verdandiviral MCPs and Topo mini-A, respectively. The trees were visualized with iTOL ^85^.

## Data availability

The raw reads and genome sequences from the metagenomes described in this study are available at NCBI under BioProject PRJNAXXXXX. Viral and MGE sequences are available at NCBI under BioProject PRJNAXXXXX.

## Acknowledgements

We thank the crews, technical staff and shipboard scientists of the DV Chikyu for the operation and sampling during CK06-06 cruise in 2006. We are grateful to Miho Hirai and Yoshihiro Takaki for the library construction and sequencing of the subseafloor samples from off Shimokita. The work in the M.K. laboratory is supported by grants from the l’Agence Nationale de la Recherche (ANR-20-CE20-0009-02) and Ville de Paris (Emergence(s) project MEMREMA). S.M. was partly supported by the Metchnikov fellowship from Campus France. N.Y. and E.V.K. are supported by the Intramural Research Program of the National Institutes of Health of the USA (National Library of Medicine). The work by C.R and J. S. is funded by the Australian Research Council (ARC) Future Fellow Award (FT170100213) awarded to CR. T.N. was partly supported by JSPS KAKENHI Grant Number JP19H05684 within JP19H05679 (Post-Koch Ecology).

## SUPPLEMENTARY INFORMATION

### Supplementary Figures

**Figure S1.**
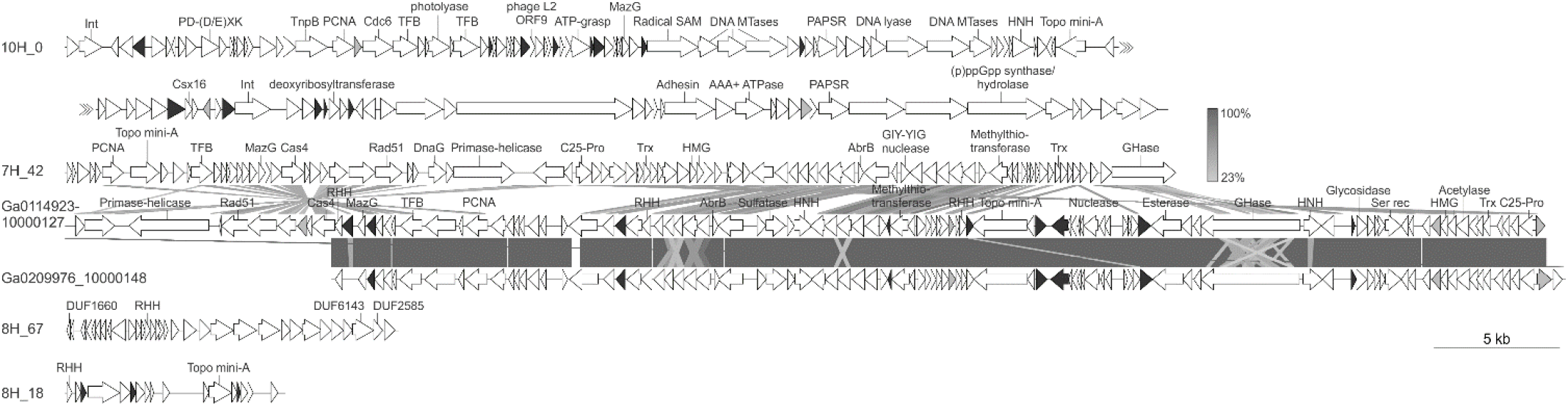
Genome maps of Asgard archaeal CRISPR-targeted MGEs. Genes encoding putative DNA-binding proteins with Zn-binding and helix-turn-helix domains are colored in black and grey, respectively. Grey shading connects genes displaying sequence similarity at the protein level, with the percent of sequence identity depicted with different shades of grey Abbreviations: Int, integrase; PD-(D/E)XK, PD-(D/E)XK family nuclease; PCNA, proliferating cellular nuclear antigen; TFB, transcription initiation factor B; MTase, methyltransferase; HNH, HNH family nuclease; PAPSR, phosphoadenosine phosphosulfate reductase; C25-Pro, C25-family protease; Trx, thioredoxin; GHase, glycoside hydrolase; RHH, ribbon-helix-helix domain-containing DNA binding protein; Ser rec, serine superfamily recombinase; HMG, high mobility group domain-containing chromatin-associated protein.

**Figure S2.**
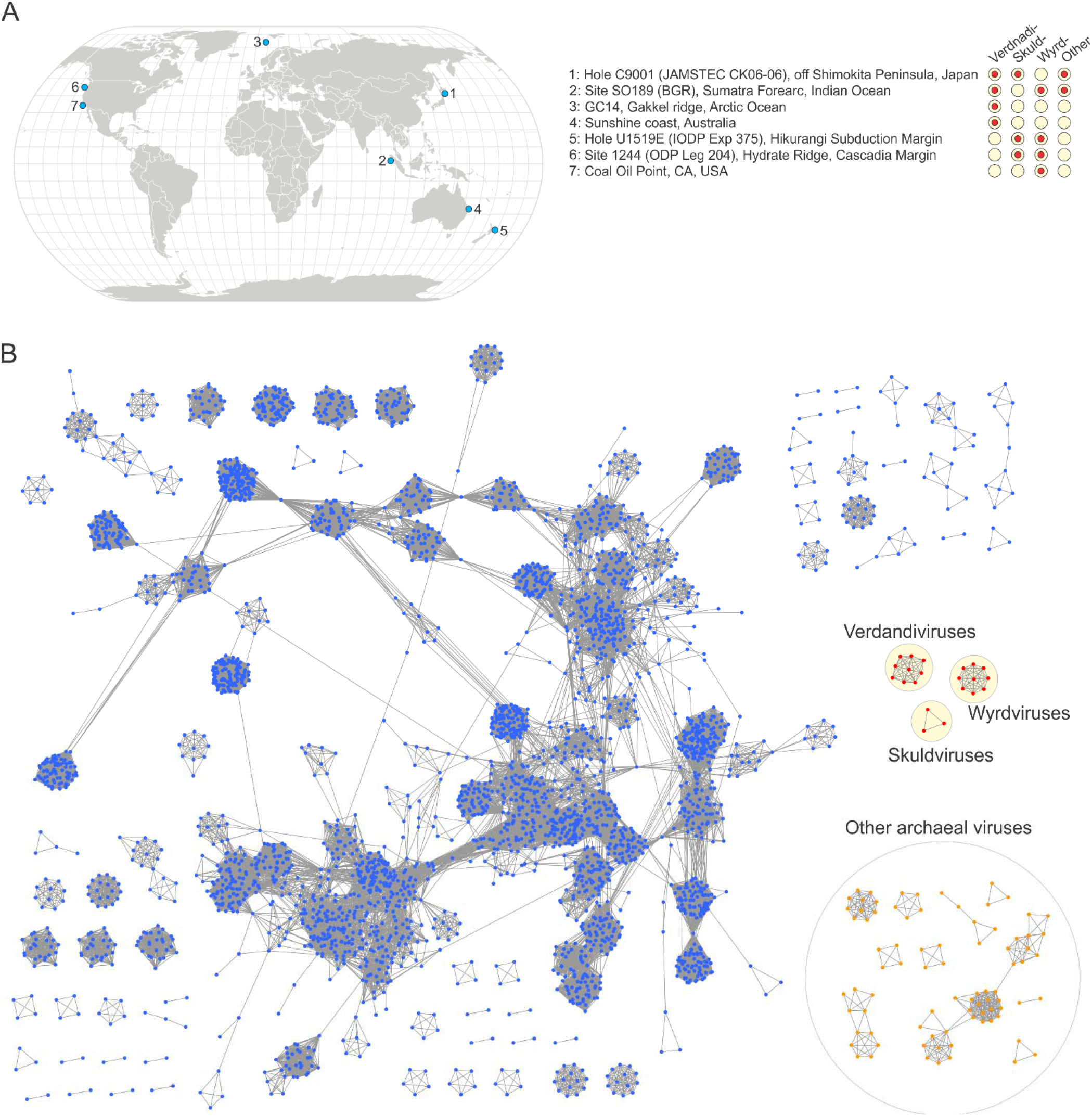
Asgard archaeal viruses and MGEs. A. Geographical distribution of Asgard archaeal viruses and MGEs. Filled circles next to geographical locations signify the presence of the corresponding virus groups in sediment samples from that location. B. The network-based analysis of shared protein clusters (PCs) among Asgard archaeal viruses and the prokaryotic dsDNA viruses. The nodes represent viral genomes, and the edges represent the strength of connectivity between each genome based on shared PCs. Nodes representing genomes of the three groups of Asgard archaeal viruses are in shown in red and the three groups are circled with yellow background. Nodes corresponding to other bacterial and archaeal viruses are shown in blue and orange, respectively.

**Figure S3.**
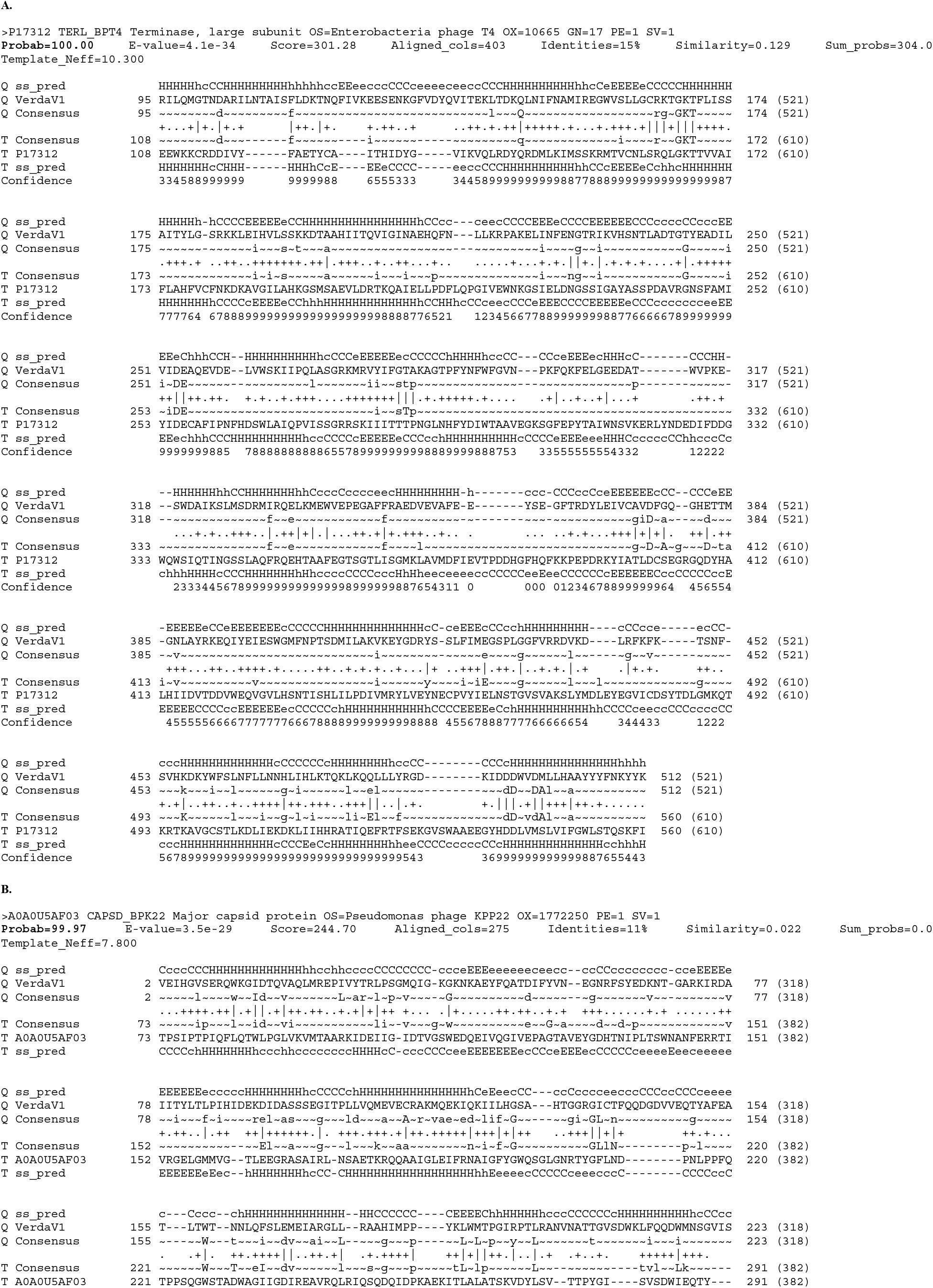
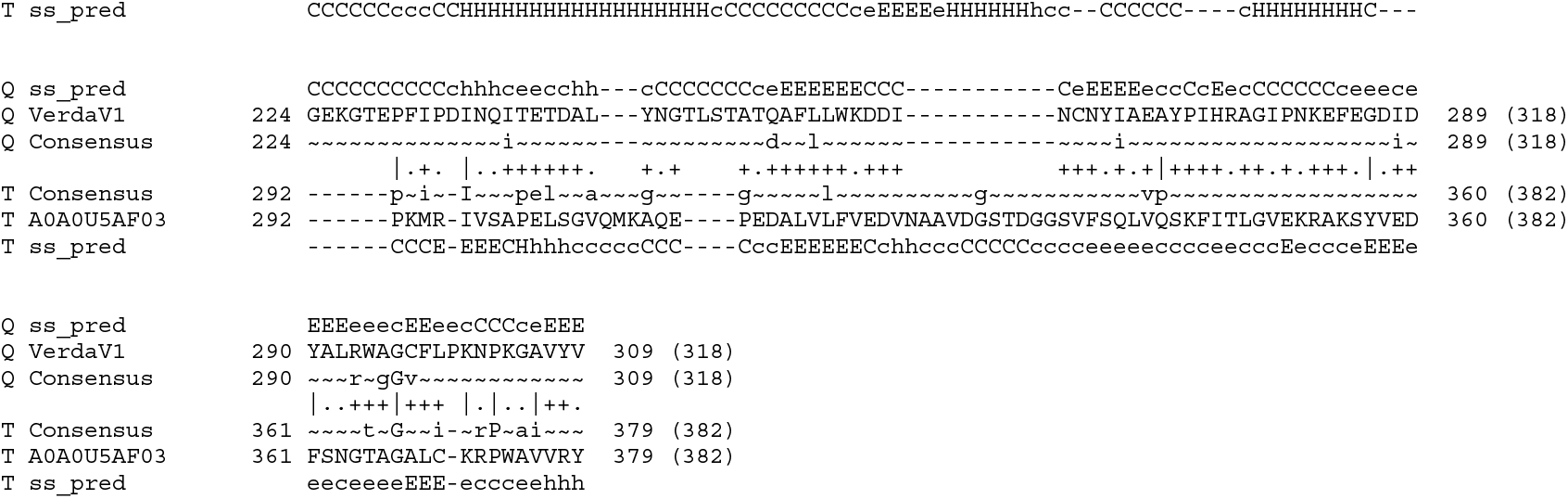
Results of the HHsearch analysis seeded with the putative large terminase subunit (A) and the major capsid proteins (B) of verdandivirus VerdaV1. H(h), α-helix; E(e), β-strand; C(c), coil.

**Figure S4.**
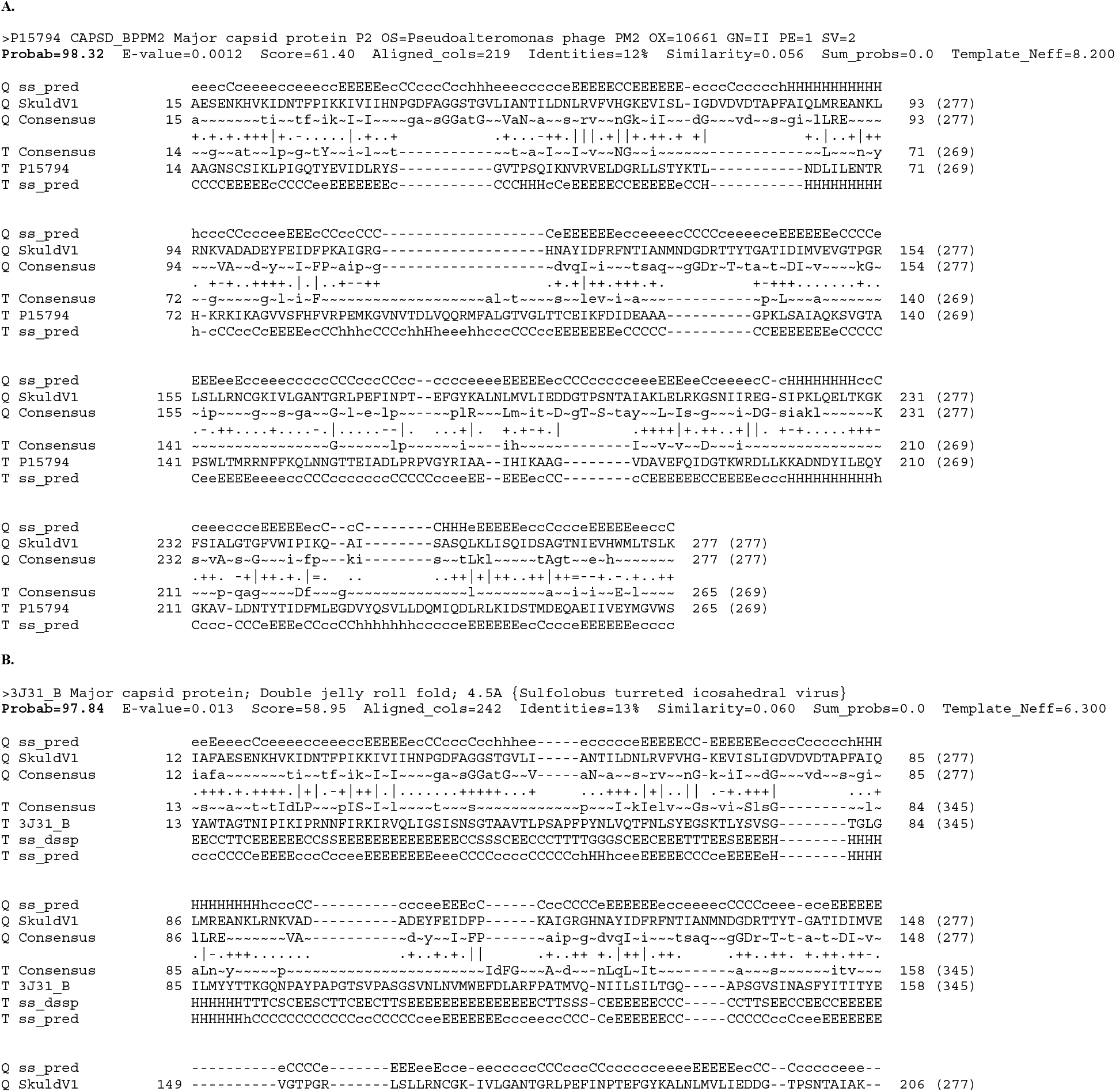
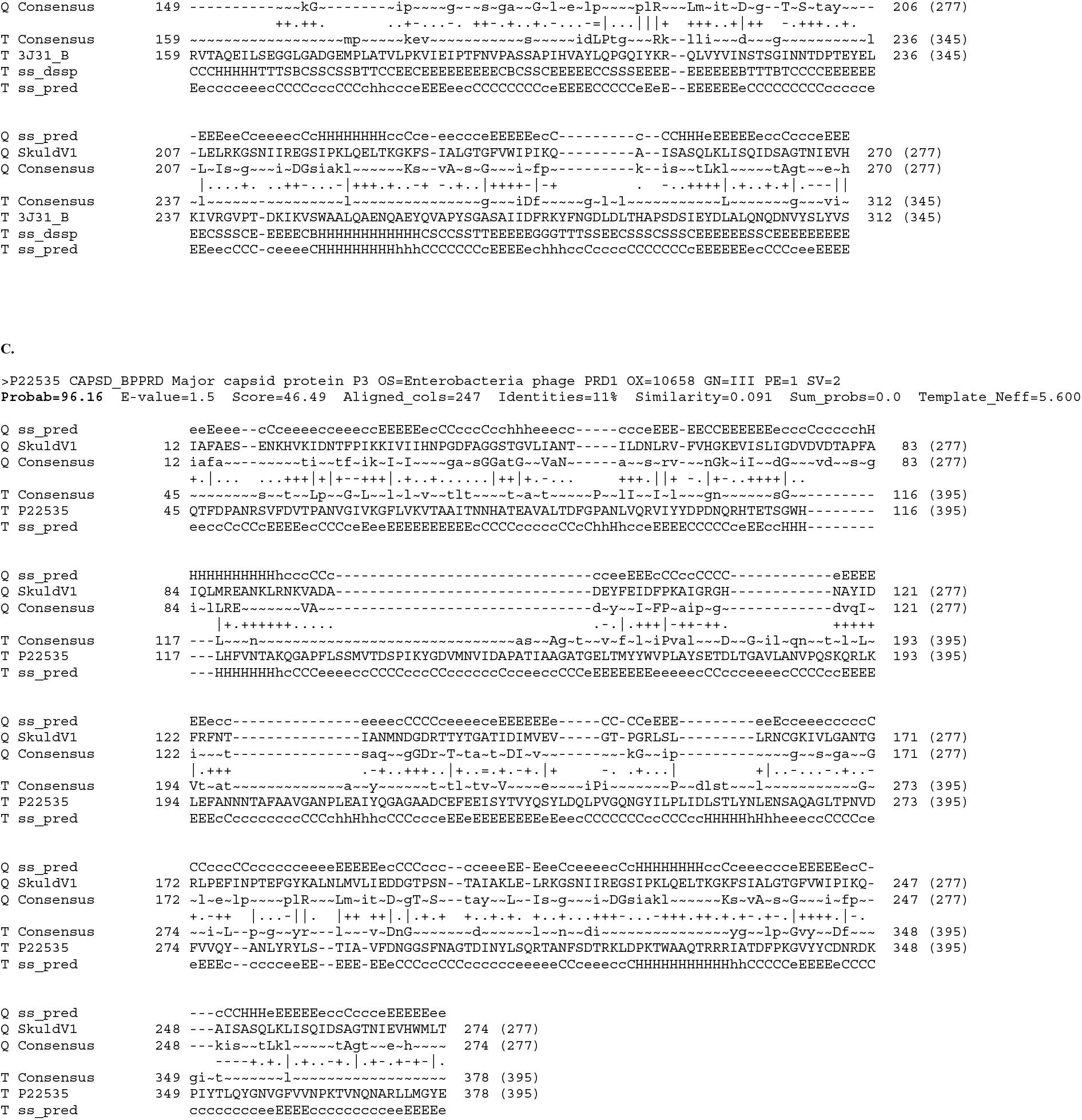
HHsearch profile-profile comparisons between the putative major capsid protein of skuldvirus SkuldV1 and the double jelly-roll major capsid proteins of (A) corticovirus PM2, (B) turrivirus STIV, and (C) tectivirus PRD1. H(h), α-helix; E(e), β-strand; C(c), coil.

**Figure S5.**
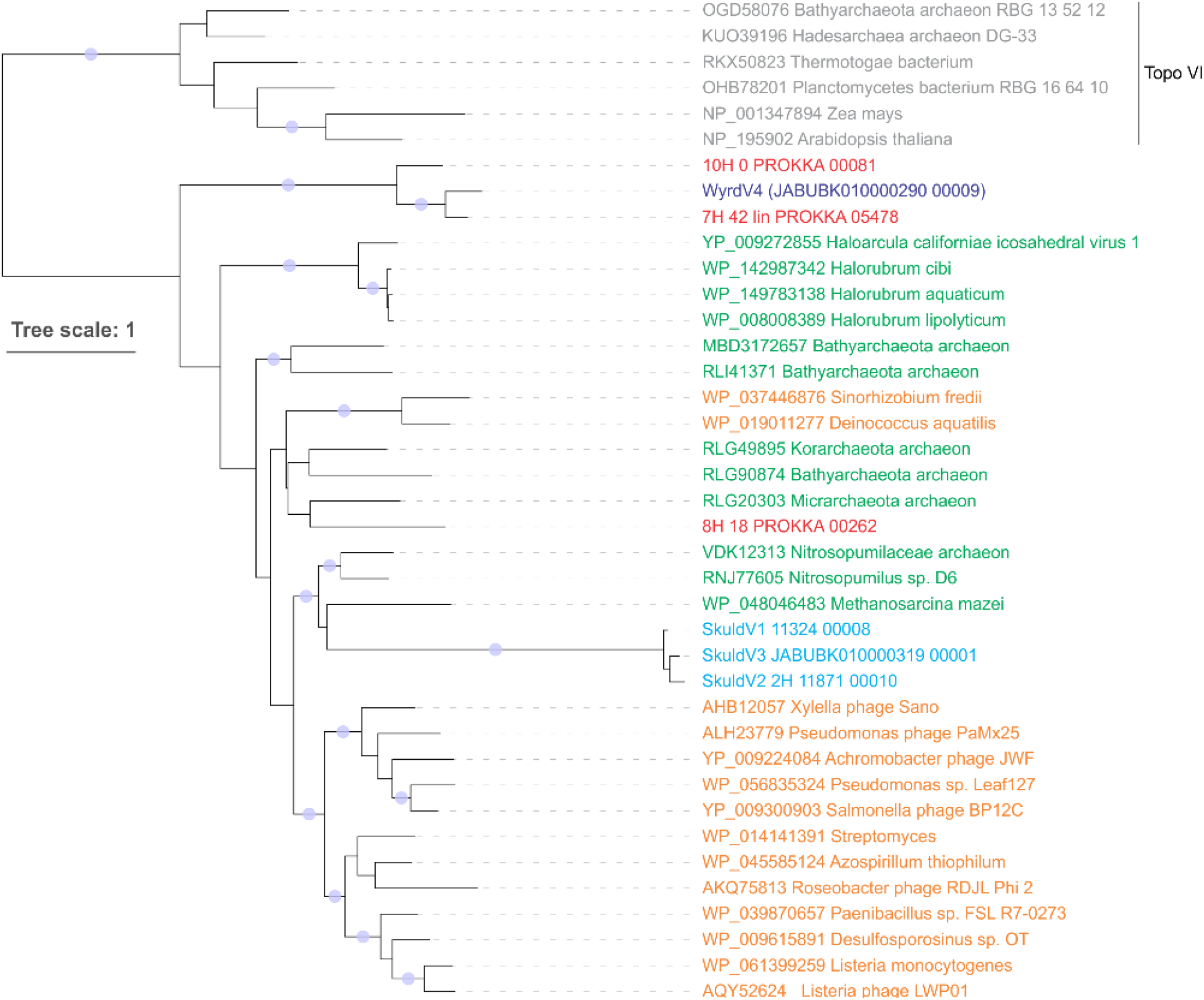
Maximum likelihood phylogenetic tree of Topo mini-A proteins. Proteins of skuldviruses and wyrdviruses are shown in cyan and dark blue, respectively, whereas those encoded by other Asgard archaeal MGE are shown n red. Other Topo mini-A homologs encoded by archaea and bacteria (or their corresponding viruses) are shown in green and orange, respectively. Topo VI proteins were used as an outgroup and are colored grey. The tree was constructed using the automatic optimal model selection (LG+R5). The scale bar represents the number of substitution per site. Circles at the nodes denote aLRT SH-like branch support values larger than 90%.

**Figure S6.**
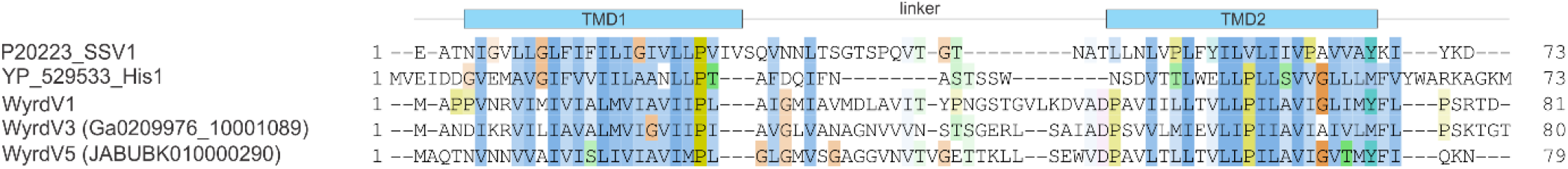
Sequence alignment of the major capsid proteins of selected wyrdviruses, fusellovirus SSV1 and halspivirus His1. TMD, transmembrane domain.

### Supplementary tables

**Table S1.** Basic statistics of Asgardarchaeota MAGs sequenced in this study (Table S1.xlsx).

**Table S2.**
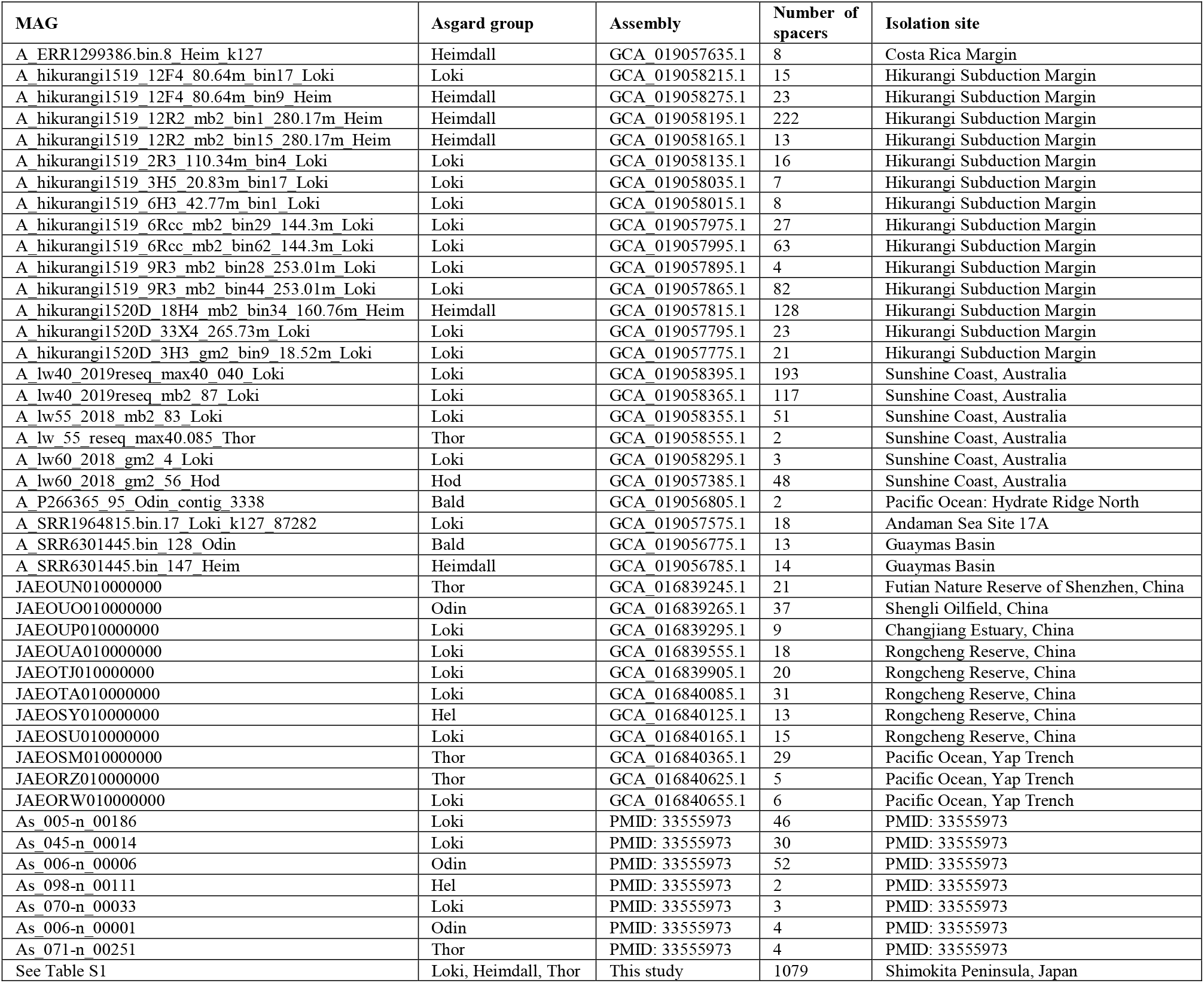
Asgardarchaeota MAGs containing CRISPR arrays.

**Table S3.**
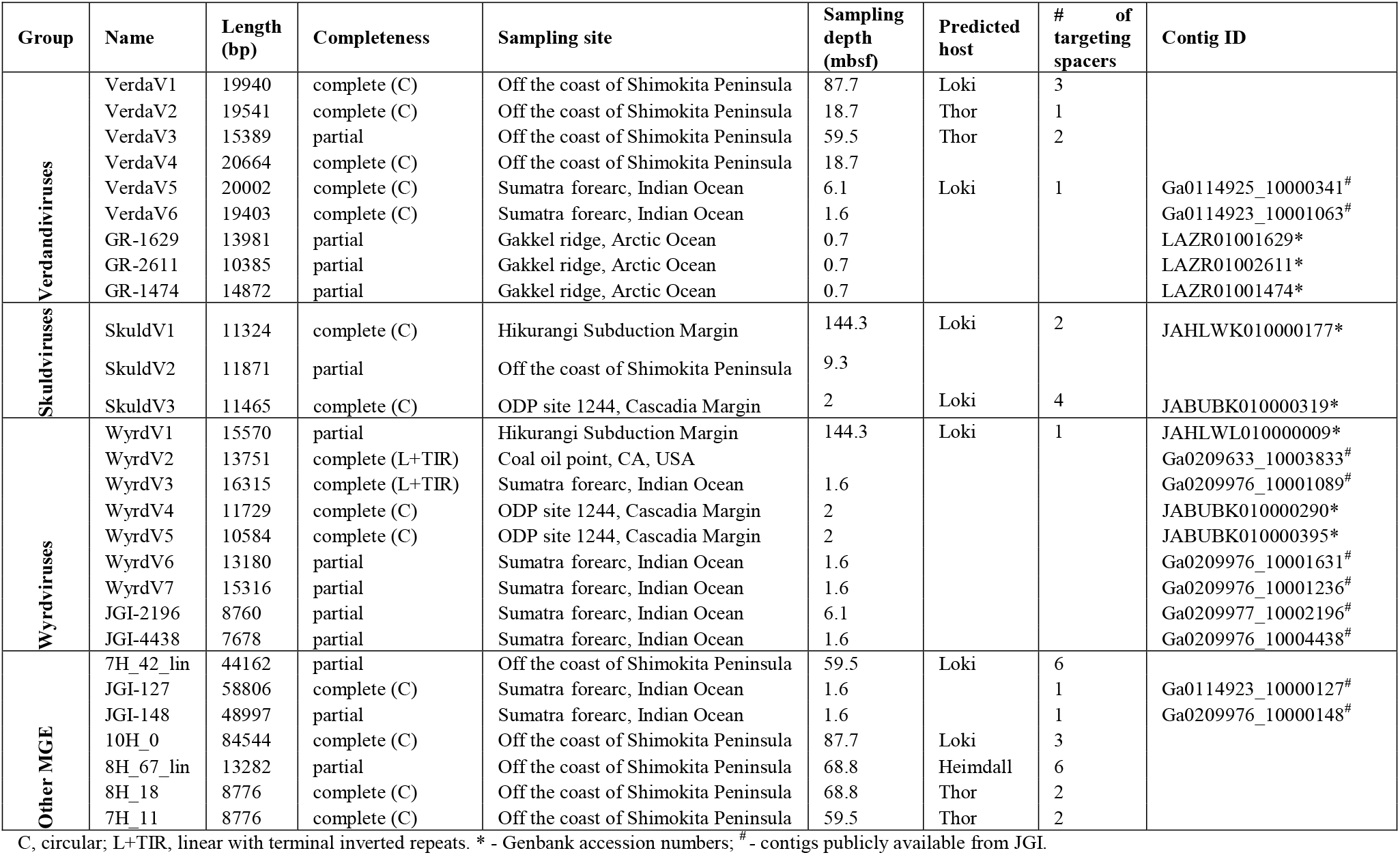
Asgard archaeal viruses and other MGEs.

**Table S4.** Spacer targeting of the putative Asgard archaeal viruses and MGEs (see Table S4.xlsx).

**Table S5.** Annotations of proteins encoded by Asgard archaeal viruses and MGEs (see Table S5.xlsx).

## References

1 Liu, Y. et al. Expanded diversity of Asgard archaea and their relationships with eukaryotes. Nature 593, 553–557 (2021).

2 Dombrowski, N., Teske, A. P. & Baker, B. J. Expansive microbial metabolic versatility and biodiversity in dynamic Guaymas Basin hydrothermal sediments. Nat Commun 9, 4999 (2018).

3 Wong, H. L. et al. Disentangling the drivers of functional complexity at the metagenomic level in Shark Bay microbial mat microbiomes. ISME J 12, 2619–2639 (2018).

4 Liu, Y. et al. Comparative genomic inference suggests mixotrophic lifestyle for Thorarchaeota. ISME J 12, 1021–1031 (2018).

5 Zaremba-Niedzwiedzka, K. et al. Asgard archaea illuminate the origin of eukaryotic cellular complexity. Nature 541, 353–358 (2017).

6 Seitz, K. W., Lazar, C. S., Hinrichs, K. U., Teske, A. P. & Baker, B. J. Genomic reconstruction of a novel, deeply branched sediment archaeal phylum with pathways for acetogenesis and sulfur reduction. ISME J 10, 1696–705 (2016).

7 Spang, A. et al. Complex archaea that bridge the gap between prokaryotes and eukaryotes. Nature 521, 173–179 (2015).

8 Sun, J. et al. Recoding of stop codons expands the metabolic potential of two novel Asgardarchaeota lineages. ISME Commun 1, 30 (2021).

9 Seitz, K. W. et al. Asgard archaea capable of anaerobic hydrocarbon cycling. Nat Commun 10, 1822 (2019).

10 Farag, I. F., Zhao, R. & Biddle, J. F. “Sifarchaeota,” a novel Asgard phylum from Costa Rican sediment capable of polysaccharide degradation and anaerobic methylotrophy. Appl Environ Microbiol 87, e02584–20 (2021).

11 Zhang, J. W. et al. Newly discovered Asgard archaea Hermodarchaeota potentially degrade alkanes and aromatics via alkyl/benzyl-succinate synthase and benzoyl-CoA pathway. ISME J 15, 1826–1843 (2021).

12 Cai, M. et al. Diverse Asgard archaea including the novel phylum Gerdarchaeota participate in organic matter degradation. Sci China Life Sci 63, 886–897 (2020).

13 Rinke, C. et al. A standardized archaeal taxonomy for the Genome Taxonomy Database. Nat Microbiol 6, 946–959 (2021).

14 Imachi, H. et al. Isolation of an archaeon at the prokaryote-eukaryote interface. Nature 577, 519–525 (2020).

15 Lopez-Garcia, P. & Moreira, D. The Syntrophy hypothesis for the origin of eukaryotes revisited. Nat Microbiol 5, 655–667 (2020).

16 Da Cunha, V., Gaia, M., Nasir, A. & Forterre, P. Asgard archaea do not close the debate about the universal tree of life topology. PLoS Genet 14, e1007215 (2018).

17 Da Cunha, V., Gaia, M., Gadelle, D., Nasir, A. & Forterre, P. Lokiarchaea are close relatives of Euryarchaeota, not bridging the gap between prokaryotes and eukaryotes. PLoS Genet 13, e1006810 (2017).

18 Baquero, D. P. et al. Structure and assembly of archaeal viruses. Adv Virus Res 108, 127–164 (2020).

19 Prangishvili, D. et al. The enigmatic archaeal virosphere. Nat Rev Microbiol 15, 724–739 (2017).

20 Dellas, N., Snyder, J. C., Bolduc, B. & Young, M. J. Archaeal Viruses: Diversity, Replication, and Structure. Annu Rev Virol 1, 399–426 (2014).

21 Makarova, K. S. et al. Unprecedented diversity of unique CRISPR-Cas-related systems and Cas1 homologs in Asgard archaea. CRISPR J 3, 156–163 (2020).

22 Makarova, K. S. et al. Evolutionary classification of CRISPR-Cas systems: a burst of class 2 and derived variants. Nat Rev Microbiol 18, 67–83 (2020).

23 Jackson, S. A. et al. CRISPR-Cas: Adapting to change. Science 356, eaal5056 (2017).

24 Coclet, C. & Roux, S. Global overview and major challenges of host prediction methods for uncultivated phages. Curr Opin Virol 49, 117–126 (2021).

25 Nunoura, T. et al. Variance and potential niche separation of microbial communities in subseafloor sediments off Shimokita Peninsula, Japan. Environ Microbiol 18, 1889–906 (2016).

26 Nurk, S., Meleshko, D., Korobeynikov, A. & Pevzner, P. A. metaSPAdes: a new versatile metagenomic assembler. Genome Res 27, 824–834 (2017).

27 Antipov, D., Raiko, M., Lapidus, A. & Pevzner, P. A. Metaviral SPAdes: assembly of viruses from metagenomic data. Bioinformatics 36, 4126–4129 (2020).

28 Glass, J. B. et al. Microbial metabolism and adaptations in Atribacteria-dominated methane hydrate sediments. bioRxiv doi: https://doi.org/10.1101/536078 (2021).

29 Dion, M. B., Oechslin, F. & Moineau, S. Phage diversity, genomics and phylogeny. Nat Rev Microbiol 18, 125–138 (2020).

30 Iranzo, J., Krupovic, M. & Koonin, E. V. The Double-Stranded DNA Virosphere as a Modular Hierarchical Network of Gene Sharing. mBio 7(2016).

31 Koonin, E. V. et al. Global Organization and Proposed Megataxonomy of the Virus World. Microbiol Mol Biol Rev 84 (2020).

32 Baek, M. et al. Accurate prediction of protein structures and interactions using a three-track neural network. Science doi: 10.1126/science.abj8754 (2021).

33 Wang, Z. et al. Structure of the Marine Siphovirus TW1: Evolution of Capsid-Stabilizing Proteins and Tail Spikes. Structure 26, 238–248 e3 (2018).

34 Hendrix, R. W. Tail length determination in double-stranded DNA bacteriophages. Curr Top Microbiol Immunol 136, 21–9 (1988).

35 Mahony, J. et al. Functional and structural dissection of the tape measure protein of lactococcal phage TP901-1. Sci Rep 6, 36667 (2016).

36 Pope, W. H. et al. Whole genome comparison of a large collection of mycobacteriophages reveals a continuum of phage genetic diversity. Elife 4, e06416 (2015).

37 Krupovic, M., Forterre, P. & Bamford, D. H. Comparative analysis of the mosaic genomes of tailed archaeal viruses and proviruses suggests common themes for virion architecture and assembly with tailed viruses of bacteria. J Mol Biol 397, 144–60 (2010).

38 Gussow, A. B. et al. Machine-learning approach expands the repertoire of anti-CRISPR protein families. Nat Commun 11, 3784 (2020).

39 Li, Y. & Bondy-Denomy, J. Anti-CRISPRs go viral: The infection biology of CRISPR-Cas inhibitors. Cell Host Microbe 29, 704–714 (2021).

40 Krupovic, M. & Bamford, D. H. Virus evolution: how far does the double beta-barrel viral lineage extend? Nat Rev Microbiol 6, 941–8 (2008).

41 Hong, C. et al. A structural model of the genome packaging process in a membrane-containing double stranded DNA virus. PLoS Biol 12, e1002024 (2014).

42 Frickey, T. & Lupas, A. CLANS: a Java application for visualizing protein families based on pairwise similarity. Bioinformatics 20, 3702–4 (2004).

43 Yutin, N., Bäckström, D., Ettema, T. J. G., Krupovic, M. & Koonin, E. V. Vast diversity of prokaryotic virus genomes encoding double jelly-roll major capsid proteins uncovered by genomic and metagenomic sequence analysis. Virol J 15, 67 (2018).

44 Oksanen, H. M. & ICTV Report Consortium. ICTV Virus Taxonomy Profile: *Corticoviridae*. J Gen Virol 98, 888–889 (2017).

45 Krupovic, M. & Bamford, D. H. Putative prophages related to lytic tailless marine dsDNA phage PM2 are widespread in the genomes of aquatic bacteria. BMC Genomics 8, 236 (2007).

46 Abrescia, N. G. et al. Insights into virus evolution and membrane biogenesis from the structure of the marine lipid-containing bacteriophage PM2. Mol Cell 31, 749–61 (2008).

47 Kazlauskas, D., Varsani, A., Koonin, E. V. & Krupovic, M. Multiple origins of prokaryotic and eukaryotic single-stranded DNA viruses from bacterial and archaeal plasmids. Nat Commun 10, 3425 (2019).

48 Takahashi, T. S. et al. Expanding the type IIB DNA topoisomerase family: identification of new topoisomerase and topoisomerase-like proteins in mobile genetic elements. NAR Genom Bioinform 2, lqz021 (2020).

49 Krupovic, M., Quemin, E. R., Bamford, D. H., Forterre, P. & Prangishvili, D. Unification of the globally distributed spindle-shaped viruses of the Archaea. J Virol 88, 2354–8 (2014).

50 Bath, C., Cukalac, T., Porter, K. & Dyall-Smith, M. L. His1 and His2 are distantly related, spindle-shaped haloviruses belonging to the novel virus group, Salterprovirus. Virology 350, 228–39 (2006).

51 Hong, C. et al. Lemon-shaped halo archaeal virus His1 with uniform tail but variable capsid structure. Proc Natl Acad Sci U S A 112, 2449–54 (2015).

52 Quemin, E. R. et al. Sulfolobus Spindle-Shaped Virus 1 Contains Glycosylated Capsid Proteins, a Cellular Chromatin Protein, and Host-Derived Lipids. J Virol 89, 11681–91 (2015).

53 Roux, S. et al. Cryptic inoviruses revealed as pervasive in bacteria and archaea across Earth’s biomes. Nat Microbiol 4, 1895–1906 (2019).

54 Straus, S. K. & Bo, H. E. Filamentous bacteriophage proteins and assembly. Subcell Biochem 88, 261–279 (2018).

55 Kim, J. G. et al. Spindle-shaped viruses infect marine ammonia-oxidizing thaumarchaea. Proc Natl Acad Sci U S A 116, 15645–15650 (2019).

56 Quemin, E. R. et al. Eukaryotic-like virus budding in Archaea. mBio 7, e01439–16 (2016).

57 Rahlff, J. et al. Genome-informed microscopy reveals infections of uncultivated carbon-fixing archaea by lytic viruses in Earth’s crust. bioRxiv https://doi.org/10.1101/2020.07.22.215848 (2020).

58 Bamford, D. H. et al. ICTV Virus Taxonomy Profile: *Pleolipoviridae*. J Gen Virol 98, 2916–2917 (2017).

59 Mizuno, C. M. et al. Novel haloarchaeal viruses from Lake Retba infecting Haloferax and Halorubrum species. Environ Microbiol 21, 2129–2147 (2019).

60 Raymann, K., Forterre, P., Brochier-Armanet, C. & Gribaldo, S. Global phylogenomic analysis disentangles the complex evolutionary history of DNA replication in archaea. Genome Biol Evol 6, 192–212 (2014).

61 Baquero, D. P. et al. New virus isolates from Italian hydrothermal environments underscore the biogeographic pattern in archaeal virus communities. ISME J 14, 1821–1833 (2020).

62 Gabler, F. et al. Protein Sequence Analysis Using the MPI Bioinformatics Toolkit. Curr Protoc Bioinformatics 72, e108 (2020).

63 Mara, P. et al. Viral elements and their potential influence on microbial processes along the permanently stratified Cariaco Basin redoxcline. ISME J 14, 3079–3092 (2020).

64 Summer, E. J., Gill, J. J., Upton, C., Gonzalez, C. F. & Young, R. Role of phages in the pathogenesis of Burkholderia, or ‘Where are the toxin genes in Burkholderia phages?’. Curr Opin Microbiol 10, 410–7 (2007).

65 Farlow, J. et al. Genomic characterization of three novel Basilisk-like phages infecting Bacillus anthracis. BMC Genomics 19, 685 (2018).

66 Gross, M., Marianovsky, I. & Glaser, G. MazG *- a regulator of programmed cell death in Escherichia coli. Mol Microbiol 59, 590–601 (2006).

67 Sullivan, M. B. et al. Genomic analysis of oceanic cyanobacterial myoviruses compared with T4-like myoviruses from diverse hosts and environments. Environ Microbiol 12, 3035–56 (2010).

68 Rihtman, B. et al. Cyanophage MazG is a pyrophosphohydrolase but unable to hydrolyse magic spot nucleotides. Environ Microbiol Rep 11, 448–455 (2019).

69 Krupovic, M., Cvirkaite-Krupovic, V., Iranzo, J., Prangishvili, D. & Koonin, E. V. Viruses of archaea: Structural, functional, environmental and evolutionary genomics. Virus Res 244, 181–193 (2018).

70 Werner, F. & Grohmann, D. Evolution of multisubunit RNA polymerases in the three domains of life. Nat Rev Microbiol 9, 85–98 (2011).

71 Bernard, S. M. et al. Structural basis of substrate specificity and regiochemistry in the MycF/TylF family of sugar O-methyltransferases. ACS Chem Biol 10, 1340–51 (2015).

72 Krupovic, M., Dolja, V. V. & Koonin, E. V. The LUCA and its complex virome. Nat Rev Microbiol 18, 661–670 (2020).

73 Cahill, J. & Young, R. Phage Lysis: Multiple Genes for Multiple Barriers. Adv Virus Res 103, 33–70 (2019).

74 Snyder, J. C. & Young, M. J. Lytic viruses infecting organisms from the three domains of life. Biochem Soc Trans 41, 309–13 (2013).

75 Krupovic, M., Daugelavicius, R. & Bamford, D. H. A novel lysis system in PM2, a lipid-containing marine double-stranded DNA bacteriophage. Mol Microbiol 64, 1635–48 (2007).

76 Danovaro, R. et al. Virus-mediated archaeal hecatomb in the deep seafloor. Sci Adv 2, e1600492 (2016).

77 Kaneko, M. et al. Insights into the methanogenic population and potential in subsurface marine sediments based on coenzyme F430 as a function-specific compound analysis. JACS Au In revision (2021).

78 Hirai, M. et al. Library construction from subnanogram DNA for pelagic sea water and deep-sea sediments. Microbes Environ 32, 336–343 (2017).

79 Hiraoka, S. et al. Microbial community and geochemical analyses of trans-trench sediments for understanding the roles of hadal environments. ISME J 14, 740–756 (2020).

80 Sieber, C. M. K. et al. Recovery of genomes from metagenomes via a dereplication, aggregation and scoring strategy. Nat Microbiol 3, 836–843 (2018).

81 Parks, D. H., Imelfort, M., Skennerton, C. T., Hugenholtz, P. & Tyson, G. W. CheckM: assessing the quality of microbial genomes recovered from isolates, single cells, and metagenomes. Genome Res 25, 1043–55 (2015).

82 Chaumeil, P. A., Mussig, A. J., Hugenholtz, P. & Parks, D. H. GTDB-Tk: a toolkit to classify genomes with the Genome Taxonomy Database. Bioinformatics (2019).

83 Price, M. N., Dehal, P. S. & Arkin, A. P. FastTree 2--approximately maximum-likelihood trees for large alignments. PLoS One 5, e9490 (2010).

84 Nguyen, L. T., Schmidt, H. A., von Haeseler, A. & Minh, B. Q. IQ-TREE: a fast and effective stochastic algorithm for estimating maximum-likelihood phylogenies. Mol Biol Evol 32, 268–74 (2015).

85 Letunic, I. & Bork, P. Interactive Tree Of Life (iTOL) v5: an online tool for phylogenetic tree display and annotation. Nucleic Acids Res 49, W293–W296 (2021).

86 Altschul, S. F., Gish, W., Miller, W., Myers, E. W. & Lipman, D. J. Basic local alignment search tool. J Mol Biol 215, 403–10 (1990).

87 Seemann, T. Prokka: rapid prokaryotic genome annotation. Bioinformatics 30, 2068–9 (2014).

88 Steinegger, M. et al. HH-suite3 for fast remote homology detection and deep protein annotation. BMC Bioinformatics 20, 473 (2019).

89 Sullivan, M. J., Petty, N. K. & Beatson, S. A. Easyfig: a genome comparison visualizer. Bioinformatics 27, 1009–10 (2011).

90 Bin Jang, H. et al. Taxonomic assignment of uncultivated prokaryotic virus genomes is enabled by gene-sharing networks. Nat Biotechnol 37, 632–639 (2019).

91 Shannon, P. et al. Cytoscape: a software environment for integrated models of biomolecular interaction networks. Genome Res 13, 2498–504 (2003).

92 Bland, C. et al. CRISPR recognition tool (CRT): a tool for automatic detection of clustered regularly interspaced palindromic repeats. BMC Bioinformatics 8, 209 (2007).

93 Tareen, A. & Kinney, J. B. Logomaker: beautiful sequence logos in Python. Bioinformatics 36, 2272–2274 (2020).

94 Katoh, K., Rozewicki, J. & Yamada, K. D. MAFFT online service: multiple sequence alignment, interactive sequence choice and visualization. Brief Bioinform 20, 1160–1166 (2019).

95 Capella-Gutierrez, S., Silla-Martinez, J. M. & Gabaldon, T. trimAl: a tool for automated alignment trimming in large-scale phylogenetic analyses. Bioinformatics 25, 1972–3 (2009).

